# Goal-seeking compresses neural codes for space in the human hippocampus and orbitofrontal cortex

**DOI:** 10.1101/2023.01.12.523762

**Authors:** PS Muhle-Karbe, H Sheahan, G Pezzulo, H Spiers, S Chien, NW Schuck, C Summerfield

## Abstract

Humans can navigate flexibly to meet their goals. Here, we asked how the neural representation of allocentric space is distorted by goal-directed behaviour. Participants navigated an agent to two successive goal locations in a grid world environment comprising four interlinked rooms, with a contextual cue indicating the conditional dependence of one goal location on another. Examining the neural geometry by which room and context were encoded in fMRI signals, we found that map-like representations of the environment emerged in both hippocampus and neocortex. Cognitive maps in hippocampus and orbitofrontal cortices were compressed so that locations cued as goals were coded together in neural state space, and these distortions predicted successful learning. This effect was captured by a computational model in which current and prospective locations are jointly encoded in a place code, providing a theory of how goals warp the neural representation of space in macroscopic neural signals.

## Introduction

Humans and other primates can use context to guide their decisions. During *instantaneous* choices, where stimuli evoke independent action-outcome mappings, contextual cues modulate the neural encoding of information in both sensory neocortex [1–3] and higher regions such as prefrontal cortex [4–8]. However, context can also influence *sequential* choices, where outcomes depend on a series of transitions between states and actions – for example, when navigating to a spatial goal. Many species can navigate flexibly to distinct goals based on contextual information that is unobservable or maintained in memory [9,10]. Humans can pursue different goals depending on the context: for example, a person might find their way to the local hairdresser or post office depending on the purpose of an errand. Flexible, context-dependent navigation requires that space and goals are encoded in ways that avoid mutual interference, allowing the correct destination to be reached given the context. Here, we studied the neural and computational mechanisms that make this possible in humans.

Recordings from the rodent hippocampus and connected structures have revealed much about the neural representation of allocentric space [11]. In the hippocampus, ‘place cells’ code for the current location of the animal via spatially localised firing patterns called ‘place fields’. These collectively form an internal map of the local environment, with each cell firing at a slightly different spatial location [12]. Neural codes for space have also been identified using single-cell recordings in other species, including humans [13,14], and gross spatial location can be read out from fMRI signals [15–18]. In rodents, changes in context can lead place cells to form new fields in different locations, a phenomenon known as remapping. Global (and partial) remapping have been observed after physical changes to the local environment, such as the introduction of novel colours, textures or odours [19] or the repositioning of the testing apparatus in a new room [20]. However, remapping can also occur when the context is denoted by an unobservable variable, such as a latent task rule [21], a noisy inference about the environment-generating process [22], or a prospective pathway or destination [23–25]. Sometimes, remapping may occur along a single dimension aligned with the gain of neural activity, which is called rate remapping [26].

The context provided by a spatial goal can also distort the representation of space without provoking gross changes in the neural code. Place cells tend to over-represent behaviourally significant spatial locations, and can accumulate around [27–29] or fire excess spikes at [30] a rewarded position. Place cells may also encode information about prospective as well as current locations on the spatial trajectory [31–33], and information about future states has also been observed in both hippocampal BOLD signals [34,35] and intracranial recordings from the human medial temporal lobe (MTL) [13,14,36]. It has also been claimed that spatial goals may be directly coded in the hippocampal formation [9,37]. One recent study has reported a small but dedicated population of CA1 neurons whose activity covaries with the location of a rewarding stimulus. When changes to the environment cause global remapping, these cells show the same preserved activity pattern linked to reward proximity [38]. These could be “goal” cells, a putative class of neuron that codes directly for a location that the animal seeks to reach, rather than current spatial location [39]. Recent recordings from the rodent orbitofrontal cortex provide parallel evidence of goal coding where the future goal location is present in the neural population before the animal begins its navigation to the goal [40].

Decoding a mixture of place and goal locations could produce a spatial representation that is warped by the animal’s intended destination, with regions of space containing two prospective goal locations being coded with a more similar neural code, and thus appearing closer together in the internal spatial map (we call this “goal-based spatial compression”). Here, we report that while different neocortical areas encode goals and locations in heterogenous ways, strong goal-based spatial compression is observed in the BOLD signal recorded from the human hippocampus and orbitofrontal cortex.

## Results

Human participants (n = 27) performed a spatial navigation task that involved controlling an avatar as it moved through a partially observable grid world composed of four discrete and interconnected rooms (**Fig. 1A**). Participants saw a plan (birds’ eye) view of the currently occupied room (a single 4 x 4 grid of squares); other rooms were not visible (**Fig. 1C**). One grid square in each room contained a boulder, and across the entire environment, rewards were hidden under two of the four boulders. On each trial, the avatar spawned in a random room, and participants’ task was to move it (using buttons for up, down, left, right) to collide with the two boulders that yielded rewards (with the minimum number of steps and in any order), avoiding those that were empty. Successful trial completion required both goals to be visited within a fixed time period.

**Figure 1.**
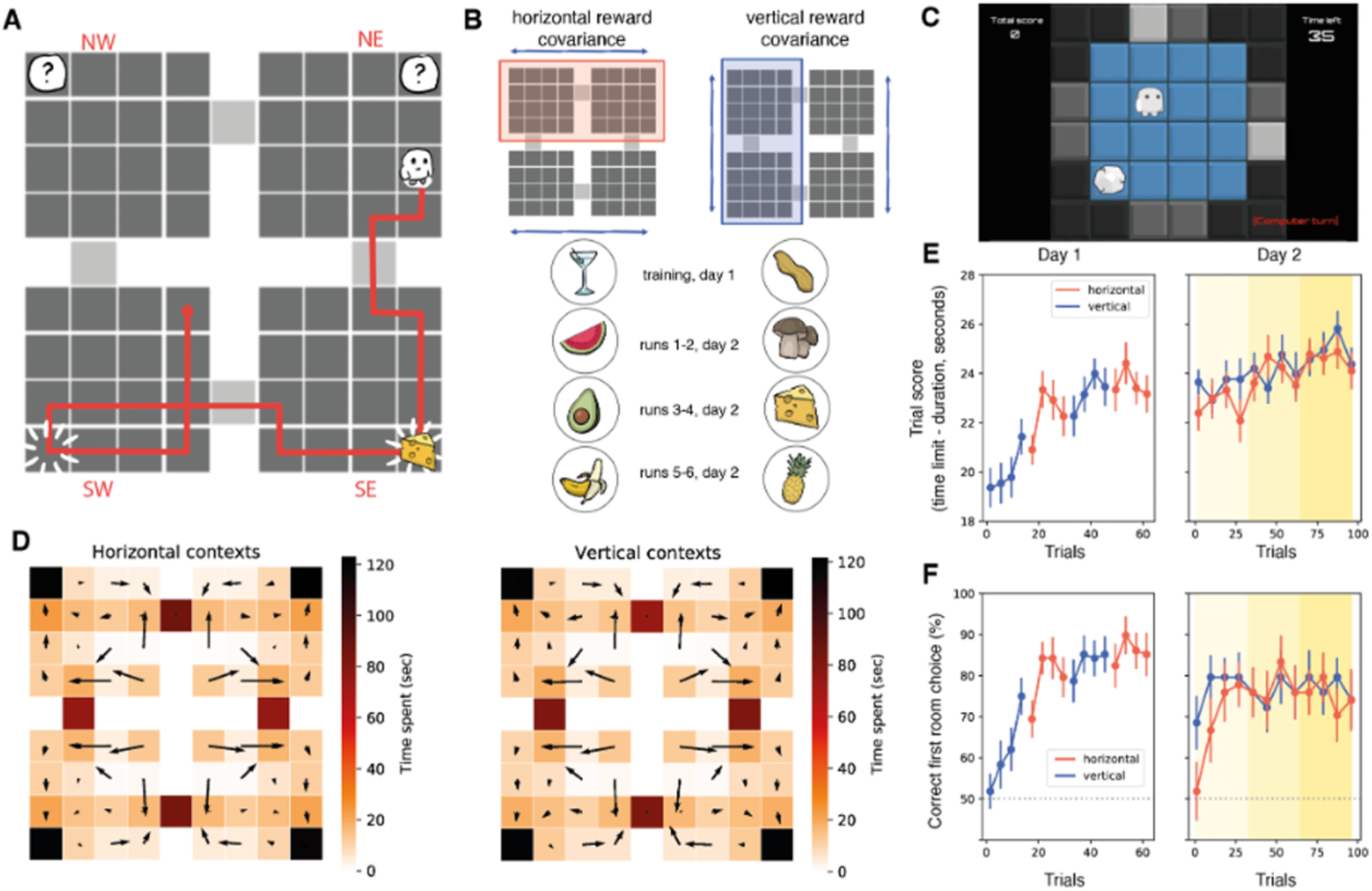
Hidden rewards covaried across four rooms. **(A)** An illustration of the four rooms environment and example reward locations under the vertical context. An example participant trajectory is shown overlaid in red. In this example, the agent starts in the south-west (SW) room, explores the start boulder and does not find a cheese reward. As cheese rewards covary vertically, the two cheese rewards must therefore be in the SE and NE rooms. **(B)** Different contexts signalled different reward covariances: one day 1 (training) martini rewards appeared in vertically adjacent rooms, while peanut rewards appeared in horizontally adjacent rooms. On day 2, three different pairs of rewards were shown, which mapped onto the same covariance structure, and the two contexts were interleaved within a run. An example ordering of runs is shown but this was balanced across participants. **(C)** Participant view of exploring in one of the rooms during training. The floors of all rooms were purple in the scanner. **(D)** Heatmaps of the average grid square occupancy per trial in each of the two contexts. Black arrows show the average transition vector from each grid square. Data are averaged across participants. Note that the average transition vectors were also well-matched when considering only those movement periods that were controlled by the participants (see **Fig. S5**). **(E)** Participant scores on each trial on days 1 (left) and 2 (right). With training, participants get faster at finding the rewards. On day 1, contexts were blocked across trials to facilitate learning, while on day 2 they were interleaved. **(F)** Participants learn to preferentially search in rooms suggested by the reward structure, and this behaviour generalises to new sets of rewards associated with each context on day 2. In panels D and E, data is shown smoothed across non-overlapping sets of 4 adjacent trials for visualisation. In panel E we show only the room choices made by human participants and exclude those made by the agent. Error bars show standard error of the mean across participants. Colour panels indicate different epochs with the same reward pairs.

At the start of each trial, participants viewed a contextual cue, which was a picture of one of two food items (**Fig.1B**). Unbeknownst to participants, each cue also revealed one reward location conditional on the other: half of the cues (‘cue H’) indicated that the rewards were in rooms lying in the same horizontal axis, while the other half (‘cue V’) indicated that the rewards were in rooms lying in the same vertical axis (with neither disclosing which specific rooms where rewards could be found). Interleaving cues from trial to trial thus ensured that a participant with perfect knowledge of the rules would on average display the same room occupancy probabilities across contexts. However, to ensure that participants also visited each room within each context, we also introduced a “robot control” phase in each trial in which participants relinquished control of the avatar to a game controller, typically moving it to a suboptimal location. This manipulation allowed us to measure BOLD signals from locations that were off the shortest path taken by expert players [41], and ensured that room occupancy probabilities and transitions were well balanced across the experiment (**Fig. 1D**).

After learning the task on an earlier day (see Methods, **Fig. 1E** and **1F** left panels), participants performed 96 trials across 6 scanner runs. In the scanner, we used three sets of physically distinct cues (food items) on runs 1-2, 3-4, and 5-6 respectively, requiring participants to generalise their knowledge of task structure across these three phases of the task (**Fig. 1B**). We plot behavioural results in **Fig. 1F**. Although participants had no way of knowing whether the first boulder they encountered (the start boulder; in the room where the avatar spawns) was rewarded or not, a participant with knowledge of the task structure can use this outcome in combination with the cue identity to exit the first room in the correct direction. Accordingly, on day 2 participants explored the start boulder on 98% of trials, and their first-choice accuracy increased rapidly across the first two runs, stabilising at about 75% (lower panels), and was significantly above chance overall (t_26_ = 8.83, p < 0.001). Time taken to complete each (correct) trial continued to decrease across the experiment (**Fig 1E** upper panels). At the end of the scanning session, participants completed a short quiz in which they were asked which room/s contained reward/s, given the presence or absence of rewards in other rooms. For example, “*You have just found a cheese in the top right room. Which room will the other cheese be in?*”. The mean score across participants was 69% ± 12% (chance performance was 34%). Quiz score positively correlated with both the average trial score on day 2 (r = 0.445, p = 0.020), and average first-choice accuracy on day 2 (r = 0.813, p < 0.001), suggesting that sequential decisions were guided by explicit knowledge about the task’s latent reward covariance structure.

To formulate neural predictions, we built a computational model that encoded the location of the avatar via simulated Gaussian place fields tiling an internal representation of the four rooms environment (**Fig. 2A**). We read out the responses elicited across the neural population as each participant moved the avatar through the four rooms, by providing empirically observed trajectories (from yoked human behavioural data) as inputs to the model (**Fig. 2B**). This allowed us to compute simulated representational dissimilarity matrices (RDMs) for each room and context (8 conditions), which were averaged before multidimensional scaling (MDS) was used to visualise their neural geometry. Without further elaboration, this model simply encodes the locations of the four rooms {Northeast (NE); Northwest (NW), Southeast (SE), Southwest (SW)} at the vertices of two perfectly parallel and aligned spatial maps, each visualised as a square plane denoting one context (**Fig. 2C**, upper left panel). This unbiased geometry was obtained with the observed human behavioural trajectories, implying that any deviations from this prediction observed in BOLD data cannot be explained by imbalance in occupancy probabilities or transition frequencies.

**Figure 2.**
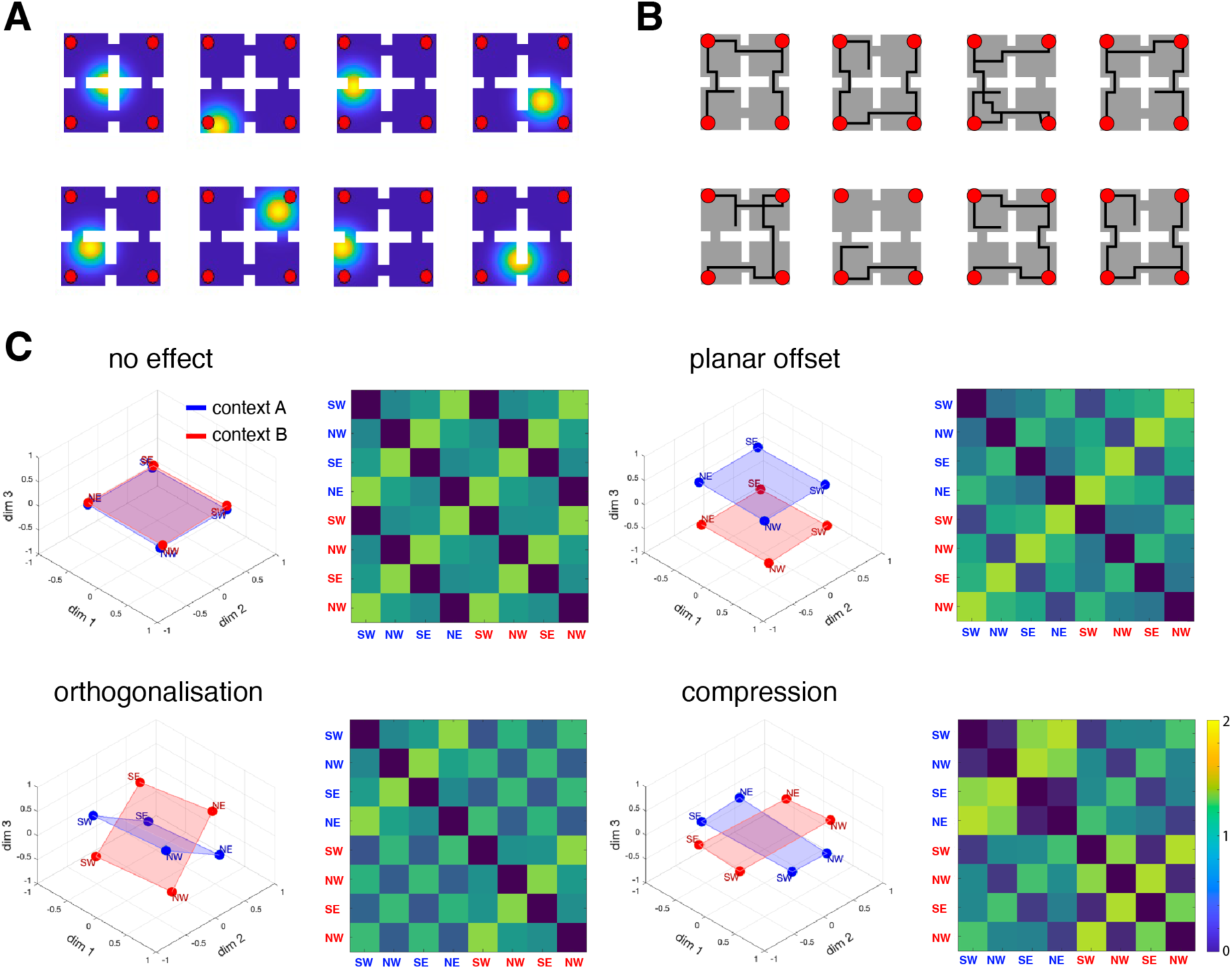
**(A)** Example simulated place fields in the model of the four rooms environment. Each panel is one neuron. White bars are walls. Red dots are boulders. The blue-yellow map shows neural tuning of a single neuron. **(B)** Example trajectories made by a participant performing the task in the scanner (black lines). Red dots are boulders. **(C)** Illustration of four representational hypotheses for different ways of separating 2D information by context. On the left of each subpanel are MDS plots. Red and blue lines and shading indicate the different contexts. The dots denote the rooms {NE, NW, SE, SW} in each context. On the right of each panel is the corresponding RDM. Colours are in units of correlation distance. The data were generated under the following parameters: *no effect*, *β* = 0, *γ* = 0.1, *ω* = 0 (a small offset is introduced for ease of visualisation); *planar separation*, *β* = 0, *γ* = 0.5, *ω* = 0; *orthogonalization*, *β* = 1, *γ* = 0, *ω* = 0; *compression*, *β* = 0, *γ* = 0, *ω* = 0.9.

However, we additionally equipped the model with three free parameters, corresponding to the hypotheses that context is encoded by (1) orthogonalization, (2) separation or (3) compression of spatial representations in each context. Firstly, we allowed the fraction of cells *β* that remapped (i.e., changed their preferred spatial location) between contexts to vary. Global remapping (*β* = 1) leads to full *orthogonalization* of the spatial map (remapping) in each context, and so it predicts that the two context-specific spatial representations (planes) should rotate to lie at 90° to one another (**Fig. 2C**, lower left panel). Secondly, we allowed a subset of cells to explicitly code for context along a dimension perpendicular to space and applied a freely varying gain factor *γ* to this neural activity, which creates a planar offset (or separation) between context representations (**Fig. 2C**, upper right panel). This model variant assumes that spatial and nonspatial variables are factorised within the neural code [10,42]. Finally, we assumed that the place cells jointly encoded the current position of the avatar and its prospective (goal) location at the goal positions on each trial, as proposed by [38]. This was achieved with a final free parameter encoding the relative mixture weight *ω* given to current and goal location. Increasing the goal weight (*ω* > 0) leads to *compression*, whereby goal locations in a shared context are neurally represented at lower distances than is warranted by their separation in physical space (i.e., the north and south rooms are closer together in the “vertical” context, and east and west in the “horizontal” context; **Fig. 2C**, lower right panel). This occurs as the neural representations of current room and future (goal) rooms are differentially mixed together in horizontal and vertical contexts. The goal weight parameter is designed to also allow the converse effect (anti-compression) when *ω* < 0, which would be consistent with other recent observations [43]. Full details of the model are provided in the Methods.

To test these hypotheses in humans, we estimated multivariate BOLD signals during navigation using a design matrix that modelled the presence of the avatar in each room (SW, NW, SE, NE) and context (H, V) during the movement period (alongside nuisance regressors) yielding an 8 x 8 RDM comparable to the model. All RDM analyses were conducted in cross-validation, comparing neural patterns between odd and even scanner runs. Goal-approach is known to be a powerful modulator of BOLD signals [41,44,45], and navigational choices are only made up until the point at which the goal room is entered, and so we begin by focussing separately on the movement phases in which participants are approaching a room containing a goal (pre-goal room period) and where they are inside a room that contains a goal (goal room period). We focus on anatomically defined neocortical regions of interest (ROIs) that have previously been implicated in goal-directed behaviour, including the prefrontal cortex (PFC), posterior parietal cortex (PPC), hippocampus (HC) and orbitofrontal cortex (OFC), as well as a control ROI in the visual cortex (shown inset in **Fig. 3C**). For each region we plotted RDMs (**Fig. 3A**) and then visualised the neural geometry in three dimensions, using multidimensional scaling (MDS; **Fig. 3B**). Finally, we complement this approach by presenting data from whole-brain searchlight analyses.

**Figure 3.**
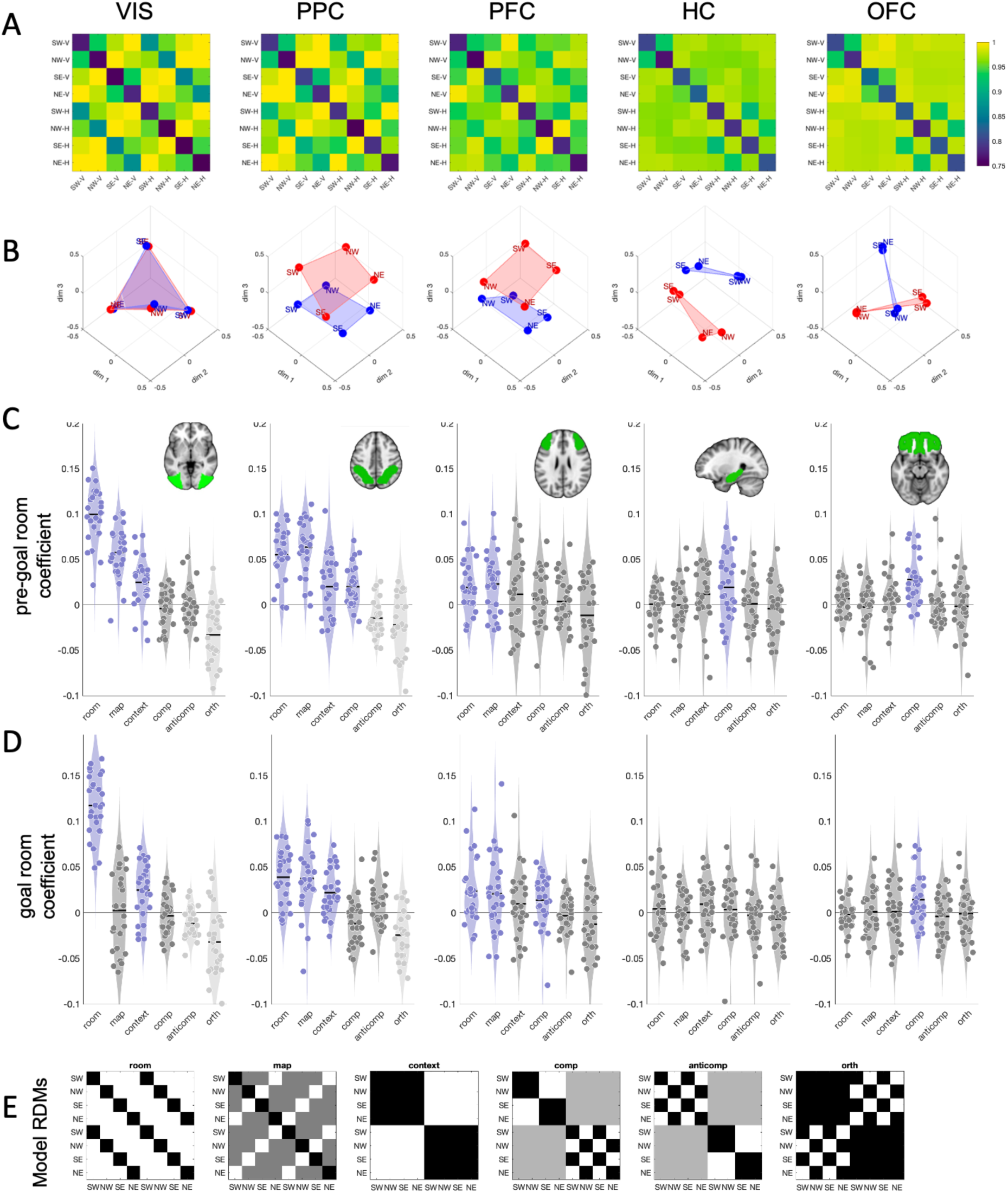
**(A)** Group average RDMs for each ROI. Each 8 × 8 RDM is ordered {SW,NW,SE,NE} for first the vertical and then the horizontal context. Warmer colours indicate greater dissimilarity, and cooler colours greater similarity. **(B)** MDS plots (from the group average RDM) for each region. Blue dots are rooms in the vertical context and red in the horizontal context. For legibility, cardinally adjacent rooms within a context are linked by lines, which collectively form a quadrilateral when allocentric space is coded in just 2 dimensions. **(C)** Violin plots showing coefficients for a competitive regression of model RDMs against each data RDM for the pre-goal room period. Each participant is an individual dot. Blue dots and shading (positive values) and light grey dots and shading (negative values) indicate p < 0.05. **(D)** Same as C but for the goal room period. **(E)** Visualisation of Model RDMs used for these analyses; lighter colours indicate greater dissimilarity.

Visual inspection of the RDMs and MDS plots revealed that BOLD signals in these ROIs coded space and context in different ways (see also supplementary videos 1-3 for a clearer visualisation of the neural geometry in 3D). Firstly, the visual cortex represented each room with a distinct neural code, but the similarity structure was only weakly related to the overall spatial layout (e.g., NE and SW rooms appeared no less similar than NE and SE rooms) and no effect of context was observed (neural manifolds for cue H and cue V conditions were aligned and superimposed; **Fig. 3B**, far left panel). In PPC and (to a lesser extent) in PFC, however, rooms were encoded at the apices of two roughly parallel quadrilateral planes, one for each context, with the rotation of the planes aligned with respect to the cardinal directions of the four rooms environment (middle left and middle panels). In other words, in PPC and PFC, allocentric space was coded with a geometry roughly matching that of the external world, even though only a single room was visible to the participant at any one time. Finally, in HC and OFC, a different pattern was observed: the neural coding of space was compressed along the irrelevant axis, so that “north” and “south” rooms are coded as adjacent in neural state space in the vertical context, and “east” and “west” are represented as neurally adjacent in the horizontal context. This compression effect was accompanied by a weak context-dependent separation, in which the two contexts were divided along another neural dimension running perpendicular (at 90°) to that encoding allocentric space (**Fig. 3B**, middle and far right panels).

To quantify these effects, we constructed model RDMs and regressed them against the neural data RDM in each ROI. We used 6 model RDMs in total and these were included competitively in the regression. The first two model RDMs related to the structure of the environment. The (i) *room* model encoded each individual room with a unique code; the (ii) *map* model encoded residual similarity structure that reflected the organisation of the four rooms into a regular quadrilateral. The remaining RDMs encoded bases for (iii) *separation* between contexts as predicted by *γ* > 0; (iv) the additional effects of both *compression* and (v) *anti-compression*, as predicted by *ω* > 0 and *ω* < 0 respectively; and (vi) the effect of *orthogonalization* as predicted by *β* > 0. The results are shown in **Fig. 3C** for the pre-goal room period and in **Fig. 3D** for the goal room period (corresponding RDMs and MDS for the latter case are shown in **Fig. S1**). The relevant statistics (corrected for multiple comparisons) are reported in **Table 1** (pre-goal room period) and **Table 2** (goal room period); we use a threshold of p < 0.01 to correct across independent ROIs. The model RDMs are also depicted in **Fig. 3E**.

**Table 1.** Statistics on regression coefficients from pre-goal room period. T-values for a test of each model RDM against zero for the data RDM from the pre-goal room period. Each row is a brain region, and each column is a predictor. Asterisks: ** p < 0.01, *** p < 0.001 after false discovery rate (FDR) correction. According to a Shapiro-Wilks test, the OFC data was not normally distributed (p = 0.03) so we additionally conducted a non-parametric (sign) test against zero; the p-value associated with this test was p < 0.001.

|  | *RDM* |  |  |  |  |  |
| --- | --- | --- | --- | --- | --- | --- |
|  | *room* | *map* | *separation* | *compress* | *anti-compress* | *orthogonalize* |
| **VIS** | 17.3*** | 10.7*** | 4.75*** | -1.04 | 0.31 | -5.44 |
| **PPC** | 9.34*** | 11.5*** | 3.06** | 4.68*** | -3.95 | -3.61 |
| **PFC** | 3.50** | 3.72*** | 1.50 | 1.13 | 0.72 | -1.44 |
| **HC** | 0.22 | -0.02 | 1.83 | 2.92** | 0.24 | -0.75 |
| **OFC** | 2.04 | -0.42 | 1.94 | 5.70*** | 0.43 | -0.24 |

**Table 2.** Statistics on regression coefficients from goal room period. T-values for a test of each model RDM coefficient against zero for the data RDM from the goal room period. Each row is a brain region, and each column is a predictor. Asterisks: ** p < 0.01, *** p < 0.001 after FDR correction.

|  | *RDM* |  |  |  |  |  |
| --- | --- | --- | --- | --- | --- | --- |
|  | *room* | *map* | *separation* | *compress* | *anti-compress* | *orthogonalize* |
| **VIS** | 18.3*** | 0.32 | 4.45*** | -0.80 | -3.69 | -4.46 |
| **PPC** | 6.77*** | 5.25*** | 4.61*** | -2.42 | 2.13 | -4.49 |
| **PFC** | 3.62*** | 2.60** | 1.43 | 2.57** | -0.81 | -1.67 |
| **HC** | 0.72 | 0.09 | 1.88 | 0.57 | -0.47 | -1.40 |
| **OFC** | -0.47 | 0.22 | 0.23 | 2.66** | -0.84 | -0.20 |

### Encoding of spatial layout

We first considered how the spatial layout of the environment was coded in BOLD signals. Our model RDMs included predictors based on unstructured room identity (*room*) and residual structure indicating the geometry of the environment (*map*). The first observation was that layout was coded most reliably in PPC and PFC. There was also a tendency for *room* to be more robustly encoded than *map* in visual cortex, especially during the goal room period (where *map* accounted for no residual variance in visual cortex) but even during the pre-goal room period *room* explained numerically more variance than *map* in visual cortex BOLD signals. This can also be seen in the MDS plots in **Fig. 3C**, where the representation of the four rooms are roughly planar in PPC but in visual cortex are folded into a tetrahedron (on a 3D simplex) to accommodate the equal similarity between rooms. Thus, there appears to be a rough progression from a more unstructured, high-dimensional representation of the spatial layout (in visual cortex) to one in which rooms lie on quadrilateral planes in neural space, mirroring the layout of the four rooms environment (in PPC).

### Goal-based spatial compression

In HC and OFC, we did not observe a neural representation of the veridical spatial layout of the environment. Rather, we saw an effect of goal-based compression, whereby rooms that were linked by virtue of shared goals were coded together. This is the pattern predicted by parameterisations of our model in which *ω* > 0, i.e., where the agent’s location and goal are jointly coded in BOLD signals. This is clearly visible in the MDS plots, where (for both HC and OFC) the north and south rooms are more proximal in the vertical context, and east and west rooms more proximal in the horizontal context (**Fig. 3C**). Compression regressors were significant in both periods for OFC and in the pre-goal room period for the HC, whereas effects of planar separation are weak or marginal in these ROIs (t-values < 2 in HC and OFC) and effects of orthogonalization are not significant. We did not observe differences in the strength of the compression effect in medial and lateral sub portions of OFC, and the pre-goal room effect was independently significant in each subregion. As a control, we also verified that these effects did not occur prior to the first goal being reached, at which point participants cannot know the optimal trajectory on that trial (**Fig. S7**).

### Model-free analyses

To complement these analyses based on a regression model, which can be difficult to interpret especially when there is partial collinearity between predictors, we adopted an approach that involved averaging selected distances (vertex pairs) across data RDMs to ask targeted questions about the neural geometry. To achieve this, we constructed “score matrices” indicating which pairs of vertices were compared with each other; these are shown in **Fig. 4A**.

**Figure 4.**
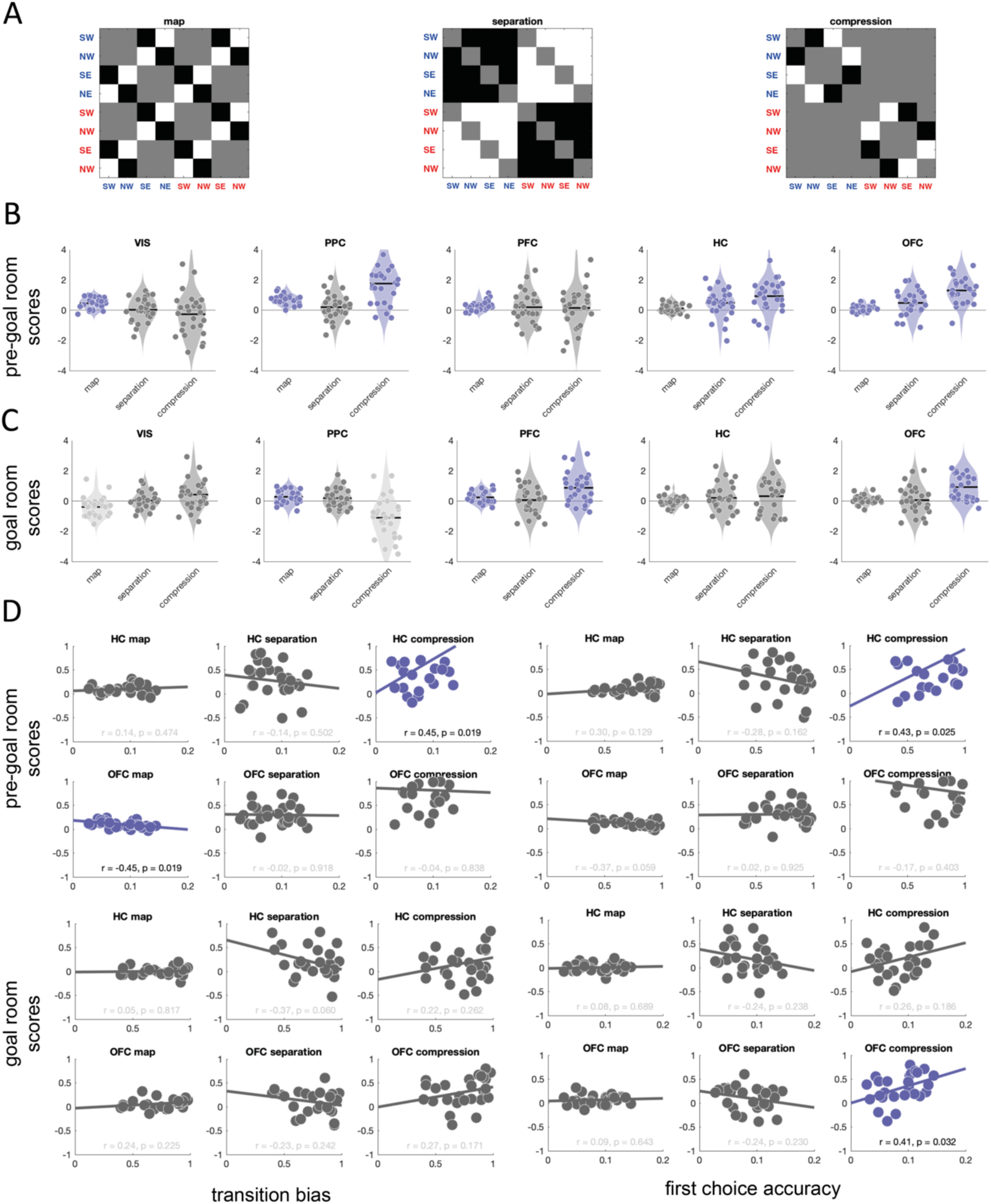
**(A)** Matrices used to compute scores. White entries are positive values (+1), black entries are negative values (-1) and grey entries are zeros (ignored). Each matrix was multiplied elementwise with the data RDM, and the resulting values summated to compute the corresponding score. The score matrices for *compression*, *separation* and *map* are fully orthogonal. **(B)** Violin plots showing scores (*map*, *separation* and *compression*) for the pre-goal room period in each region. Each dot is an individual participant. Blue dots and shading (positive values) and light grey dots and shading (negative values) indicate p < 0.05. **(C)** Same as A but for the goal room period. Below, the (C) Left: correlations between each score (see plot title) and transition bias for the HC (upper panels) and OFC (lower panels). Right: the same plots for transition bias. Blue dots denote significant (p < 0.05) correlation. **(D)** Correlations between neural scores (*map*, *separation* and *compression*) and behavioural measures (transition bias and first choice accuracy) for HC and OFC in the pre-goal room period (upper panels) and goal room period (lower panels). Each dot is a single participant, and the line is the best linear fit. Blue colouring is used to highlight significant correlations (p < 0.05).

First, we asked whether the planes for each context were roughly quadrilateral (reflecting the spatial layout of the environment). Here, we compared distances between rooms that were spatially adjacent (e.g., NE and NW) to those that were not (e.g., NE and SW), yielding a single *map score* which was zero under the null, but for which positive scores provided evidence for quadrilateral structure. In **Fig. 4B** (see also **Table 3**) we can see that there is a significant map score in all regions except HC during the pre-goal room period, and in PPC and PFC during the goal room period. Secondly, we computed a *separation score* by comparing neural distances between each room and every other room within and between contexts. Whilst the effect of separation was only marginal in the regression analysis, the separation score was reliable during the pre-goal room period for both HC and OFC. Finally, we computed a *compression score* by comparing distances between N and S and E and W rooms in each context; this score was positive if E and W rooms were neurally more proximal in the horizontal context and N and S rooms were more proximal in the vertical context, and negative for the converse.

**Table 3.** Scores for map structure, planar separation, and compression. T-values for a test of each score in each period. Each row is a brain region. Asterisks: ** p < 0.01, *** p < 0.001 after FDR correction.

|  | *Score matrix (pre-goal room)* |  |  | *Score matrix (goal room)* |  |  |
| --- | --- | --- | --- | --- | --- | --- |
|  | *Map* | *Separation* | *Compression* | *Map* | *Offset* | *Compression* |
| **VIS** | 6.56*** | 0.16 | -1.00 | -3.13 | 0.30 | 2.27 |
| **PPC** | 10.82** | 1.19 | 7.05*** | 3.44** | 1.62 | -4.33 |
| **PFC** | 4.45*** | 0.98 | 0.58 | 3.27** | 0.39 | 4.33*** |
| **HC** | 2.11 | 2.66** | 4.81*** | -0.30 | 1.18 | 1.54 |
| **OFC** | 3.01** | 3.08** | 7.84*** | 1.37 | 0.35 | 5.98*** |

The results largely mirrored those for regression analyses, with strong compression observed in HC and OFC during the pre-goal room period, and in OFC only during the goal-room period, with additional compression observed in PPC and PFC during only one of these periods. For each of these tests, statistics are shown in **Table 3**. This implies that the compression observed in HC and OFC was not an artifact of the other predictors included in our regression analysis. We also used the place cell model to create model RDMs based on the best-fitting variants of models in which each of the three parameters *β*, *γ* and *ω* were allowed to vary (or none of the three). This also confirmed that the neural data was best explained by a compression-based account in both HC and OFC (**Fig. S2**).

### Correlations between behaviour and brain activity

Next, we examined how across-cohort variation in compression scores for each brain region related to individual differences in behaviour. One way to characterise individual participant performance is *transition bias*, which is the relative fraction of transitions made horizontally and vertically between rooms in the H and V contexts (a player that understands the structure should make proportionally more horizontal transitions in H context and vertical in the V context). An alternative measure is *first choice accuracy*, which indexes whether participants’ first transition reveals that they understand the correlation structure of the spatial goals in each context. For completeness, we correlated these behavioural measures with *compression*, *separation* and *map* scores, although our main prediction was that compression would covary with performance in HC and OFC.

We observed that in the hippocampus, compression score from the pre-goal room period positively predicted both transition bias (r = 0.45, p = 0.019) and first choice accuracy (r = 0.43, p = 0.025). By contrast, in the OFC, compression score from the goal room period positively predicted first choice accuracy in the goal room period (r = 0.41, p < 0.032). We plot the results of this correlation for HC and OFC in **Fig. 4D**; results for other regions are shown in **Fig. S3**.

### Inverted neural geometry across periods

Having examined the geometries for the pre-goal room and goal-room periods, a natural next step is to explore how they relate to one another. The RDMs and corresponding MDS plots for the full period x room x context analysis are shown in **Fig. 5A-B**. In the MDS plots, the pre-goal room period is now shown in cyan (cue V) and orange (cue H), and the goal room period in blue (cue V) and red (cue H). As can be seen, the brain powerfully encodes whether the agent is currently occupying a room with a goal or not, visible as the checker pattern in the RDMs and the resulting one-dimensional offset between periods that lies along a neural dimension perpendicular to that coding allocentric space. These results are confirmed by regressing model RDMs against the full 16 x 16 data RDM (**Fig. S4**); the effect of *period* was highly significant in each region (all t-values > 13, all p-values < 0.001), with other effects mostly mirroring those described above. Note that all analyses are conducted in cross-validation, so this is unlikely to be spuriously driven by temporal autocorrelation in BOLD signals. It is however consistent with previous reports that BOLD signals are powerfully modulated on the approach to a goal [41,44]. We can see that the effects of spatial layout (with or without compression) are thus represented in two parallel planar geometries, with a large offset coding whether the agent is currently occupying the goal room or is still navigating towards it. Interestingly, in the MDS plots for OFC and HC the orientation of the planes for contexts H and V appears flipped between the two contexts, such that the coding of space and context is inverted when it is held in memory (during the pre-goal room period) and when it is being executed.

**Figure 5.**
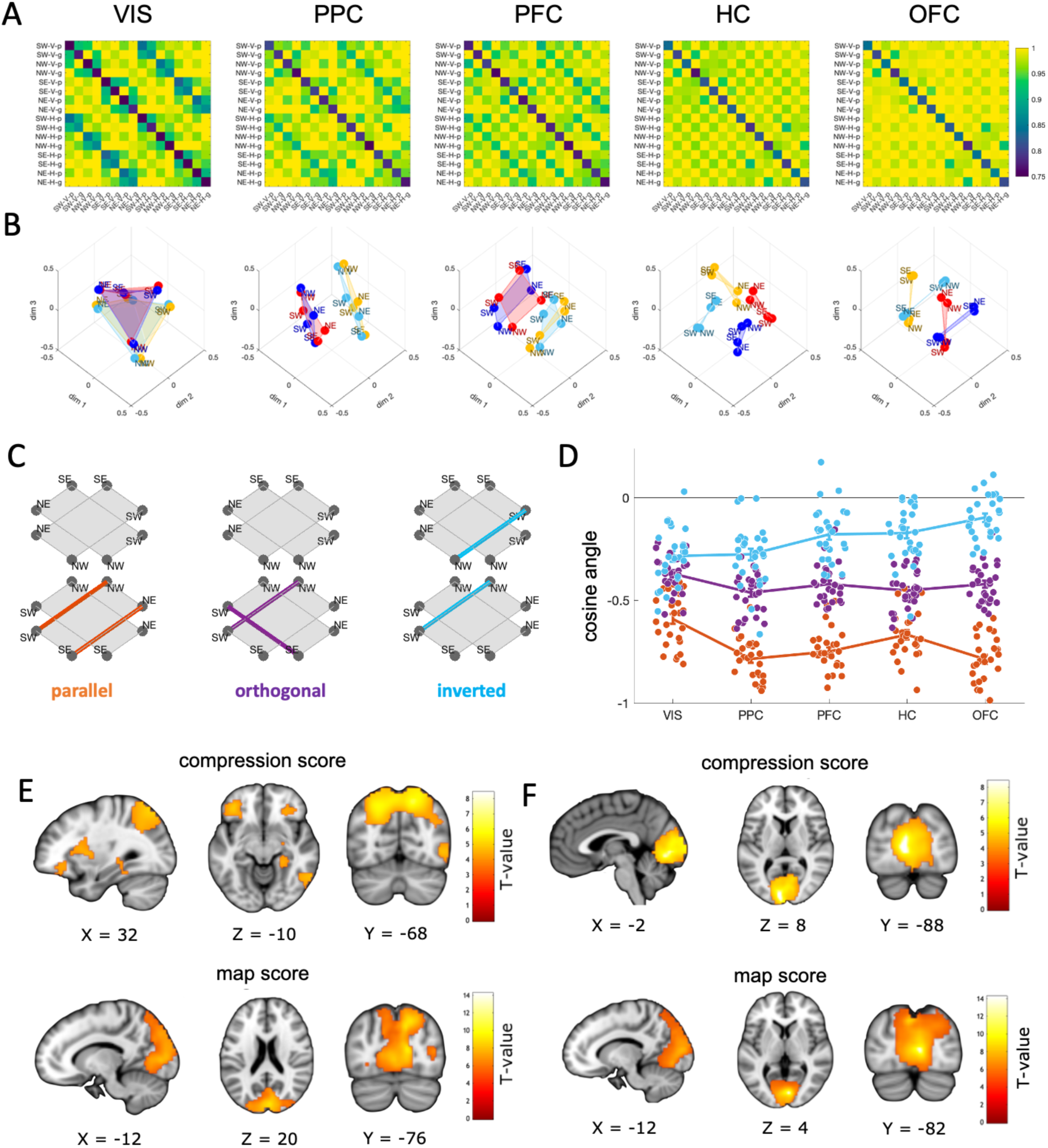
**(A)** Group average RDMs for the full period × room × context analysis in each region. Each RDM comprises the nested variables period {goal room, pre-goal room}, room {SW, NW, SE, NE} and context {vertical, horizontal}. Warmer colours signal greater dissimilarity. **(B)** MDS plots from the corresponding group average RDM. Colours denote period / context combinations: dark blue = vertical, pre-goal room; red = horizontal, pre-goal room; cyan = vertical, goal room; orange = horizontal, goal room. **(C)** Stylised model of the MDS plots in B, to illustrate those edges predicted to be parallel (left panel, orange lines), orthogonal (middle panel, purple lines) and inverted (right panel, cyan lines). **(D)** cosine angle between neural vectors for all predicted parallel, orthogonal and inverted edges, rendered onto a single plot. Dots are individual participants, and the line shows the average for each region. **(E)** Searchlight analyses: whole-brain effects of *compression score* and *map score* for the pre-goal room period, rendered onto a template brain at a threshold of p < 0.0001 uncorrected. All regions shown contain voxels significant at p < 0.05 after familywise error correction (whole-brain images thresholded with familywise error correction at p < 0.05 are shown in the supplementary materials). **(F)** Same as (E) but for goal room period.

**Table 4.** Statistics on regression coefficients from a regression modelling both pre-goal and goal room periods. T-values for a test of each model RDM coefficient against zero for the data RDM from the goal room period. Each row is a brain region, and each column is a predictor. Asterisks: ** p < 0.01, *** p < 0.001.

|  | *RDM* |  |  |  |  |  |  |
| --- | --- | --- | --- | --- | --- | --- | --- |
|  | *room* | *map* | *period* | *separation* | *compression*<br>(goal room) | *compression*<br>(pre-goal room) | *orthogonalize* |
| **VIS** | 18.35*** | 2.84** | 20.6*** | 0.60 | 2.28 | -1.00 | -4.50 |
| **PPC** | 9.52*** | 8.76*** | 20.7*** | 0.63 | -4.33 | 7.06*** | -1.39 |
| **PFC** | 3.90*** | 5.08*** | 16.7*** | 1.04 | 4.33 | 0.58 | -0.21 |
| **HC** | -0.30 | -1.47 | 13.0*** | 2.91** | 1.53 | 4.82*** | 1.90 |
| **OFC** | -1.91 | -7.46 | 14.45*** | 1.76 | 5.97*** | 7.85*** | 3.98*** |

### Cosine similarity of neural vectors within and between periods

To quantify this latter effect, we computed the angle of the high dimensional neural vector between each room/context and every other, both within periods (e.g., goal room to goal room period) and across periods (e.g., goal room to pre-goal room period). Assuming a stylised model in which the contexts were represented as compressed planes that were offset and inverted between periods (**Fig. 5C**), we averaged angles for those edges that the model predicted by this model to be parallel (e.g., common directions within a context; orange lines), orthogonal (e.g., perpendicular directions within a context; purple lines) and inverted (e.g., common directions within a context, but across periods; cyan lines). Note that this analysis was conducted in the high-dimensional space of neural activity, not its compression to 3D in the MDS plots. In **Fig. 5D** we plotted the resulting angles for each ROI, which range from fully parallel (0) to fully orthogonal (*π*/2) to inverted (*π*). The results show that across regions (but especially for HC and OFC) there is a bias for edges within a context (e.g., NE-SE) to be parallel with edges denoting a common direction in space (e.g., NW-SW); and that by contrast those edges were more orthogonal to those denoting a perpendicular direction in space (e.g., NE-NW); and that edges that denoted a common direction across periods (e.g., NE-SE in pre-goal room period and NW-SW in goal room period) had an even greater angular separation. We observed a main effect of region (F_3.6, 93.6_ = 9.91, p < 0.001) and a region × pair type interaction (F_5.9,153.0_ = 20.8, p < 0.001), with the strongest separation of angle between edge pair types in the HC and OFC relative to other regions. These results thus confirm that the neural vectors respect the geometry of the environment within each period, but are inverted between periods, especially in HC and OFC.

These analyses all rely on 5 ROIs which we chose *a priori* given their previously described involvement in context-sensitive decision-making, navigation and planning. However, to study these effects at the whole-brain level, we combined the score analysis with a whole-brain searchlight approach, allowing us to render the map, separation and compression effects onto a template brain. The results are consistent with our ROI analyses, and all regions described here contain searchlights which reach significance at the whole-brain family-wise error-corrected [FWE] level. During the pre-goal room period, we observed significant *compression score* in the posterior parietal cortex (peak t = 8.47, FWE p < 0.001), the right inferior temporal gyrus (peak [52 -63 -3], t = 4.83, FWE p = 0.003), the orbitofrontal cortex (peak [-38 36 -12], t = 4.79, FWE p = 0.005); the right putamen (peak [30 3 6], t = 5.45, FWE p = 0.024), the right hippocampus (peak [24 -39 -6], t = 5.36, FWE p = 0.032), and the right middle temporal gyrus (peak [-20 57 -18], t = 5.23 FWE p = 0.045). Significant correlations with *map score* were observed bilaterally in occipital and parietal cortices (peak [18 -84 6], t = 12.14, FWE p < 0.001), the precentral gyrus (peak [18 -84 6], t = 6.44, FWE p = 0.007), and the right insula (peak [48 3 -9], t = 6.08, FWE p = 0.015). During the goal room period, significant correlations with compression score were observed only in the medial portion of the visual cortex (peak [- 8 -90 6], t = 8.42, FWE p < 0.001), and significant correlations with map score in bilateral visual and parietal cortices (peak [18 -66 51], t = 6.55, FWE p < 0.001). No significant correlations with *separation score* were observed in either period at the chosen statistical threshold. **Fig 5F** shows a visualisation of the searchlight results for the pre-goal room period at a slightly more liberal statistical threshold (p < 0.0001, uncorrected) to facilitate illustration of smaller clusters (see **Fig. S6** for whole-brain maps FWE corrected whole-brain maps).

### Neural geometry of current and prospective locations

Finally, we asked how spatial goals were represented in each ROI, and how their geometry related to that representing the current location in space. To ensure sufficient trial counts for this analysis, we collapsed over context, and modelled the BOLD data at the first level GLM with regressors coding for the currently occupied room and the location of the current goal, in a 4 × 4 factorial design. This allowed us to construct model RDMs that encoded allocentric space as individual rooms or as a map (*room* and *map*, exactly as above) alongside new model RDMs that encoded current goal locations as individual rooms or as a map (*goalroom* and *goalmap*; **Fig. 6A**). Regression against the 16 x 16 data RDM revealed that *goalmap* was significant in visual cortex, PPC and PFC, in addition to *map*. No effects were significant in HC or OFC, presumably because collapsing over orthogonal contexts removed the relevant subspaces in which rooms and goals are represented.

**Figure 6.**
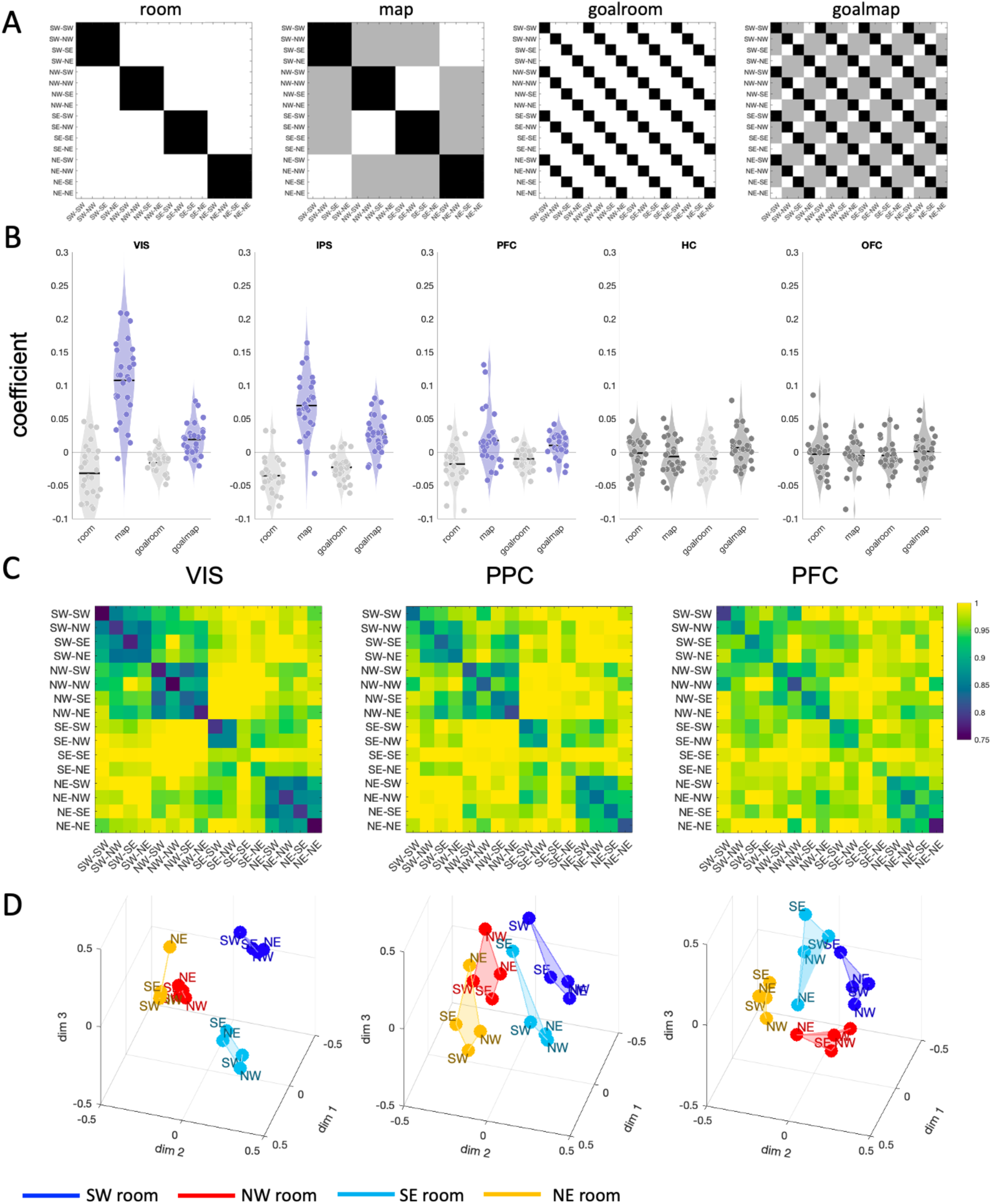
**(A)** Model RDMs used to test the neural geometry of room and goal representations. RDMs are constructed from nested goal {SW, NW, SE, NE} and room {SW, NW, SE, NE} variables. **(B)** Violin plots showing parameter estimates for each model RDM regressed competitively against the data RDM. Each dot is an individual participant. Blue dots and shading (positive values) and light grey dots and shading (negative values) indicate p < 0.05. **(C)** Group average data RDMs for visual cortex, PPC and PFC (HC and OFC showed no significant effects). **(D)** MDS plots constructed from corresponding group average RDM for each region. Colours denote room (blue = SW, red = NW, cyan = SE, orange = NE). Rooms are organised into an approximate quadrilateral, and goals (within each room) are similarly arranged approximately quadrilaterally.

The full data from the regression analysis are shown in **Fig. 6B**, and RDMs for this analysis are shown in **Fig. 6C**, and the MDS plots in **Fig. 6D**. To increase legibility, we invert the plotting convention of the previous analysis and now plot different rooms (in allocentric space) in different colours (blue = SW, red = NW, cyan = SE, orange = NE), and the labels on the plot now refer to goals (where the agent is headed). In PPC and PFC, goals are represented on 4 rough quadrilaterals, one within each room that the agent could occupy, though the representation of goals is smaller in area than the representations of room in allocentric space. There is thus a clear hierarchical representation, whereby a neural map of the current goal is represented within a neural map of the current location. In PPC, the quadrilateral is visibly elongated so that the goal room condition is represented on a common plane separated from the non-goal room conditions. The pattern in visual cortex is harder to discern. In PPC and PFC, thus, spatial goals are represented in a geometric format similar to physical space itself.

## Discussion

In the current study, participants learned that contextual cues determined the pattern of spatial goals (horizontal or vertical relationship between rewarded locations) to which an avatar should be moved in a structured “four rooms” environment [46]. One major observation from our study was that the neural geometry of BOLD signals broadly reflects the allocentric spatial layout of the environment, especially in neocortical regions such as PPC and PFC. This finding complements previous reports that spatial location (in a single context) can be decoded from BOLD signals in both hippocampus and neocortex using nonlinear pattern classification methods [15–18]. We note that this encoding of spatial layout occurred even though participants were provided with a bird’s eye (grid world) rather than a first-person view of the environment, unlike in previous studies.

Our major question was how context modulates the neural representation of allocentric space in human BOLD signals. We considered three major hypotheses. Firstly, we asked whether context would lead to remapping (or “orthogonalization”) whereby population codes for space change their tuning preferences between contexts (where the rule for finding goals changes). This is implied by previous work in rodents showing that changes to the physical nature of the environment (for example, by changing the shape of the testing environment, or varying the textures or odours encountered during navigation), or even changes in internal variables (such as thirst or hunger), can cause the spatial preferences of place cells to randomly remap [19–22,26]. One salient observation in the current report is that whilst context provoked representational changes in BOLD signals in both hippocampus and neocortex, none of these changes resembled those expected if neural codes for space randomly remap, either partially or in full. The neural geometry implied by random remapping is that spatial representations become “orthogonal” or uncorrelated. By contrast, we observed that neural manifolds representing space were highly aligned across contexts in most brain regions. This resembles the “neural structure alignment” that has recently been reported to accompany decision tasks in both humans and monkeys, whereby contexts sharing common structure are represented with parallel neural geometries, potentially because this allows a decoder trained in one context to be generalised to the other [47–52].

The second hypothesis we considered was that context is represented as an independent, nonspatial dimension in neural state space. This is implied by recent findings emphasising that multiple task-relevant variables, such as location and evidence for progress towards a goal, are multiplexed in neurons with spatial selectivity, for example in the rodent hippocampal CA1 area [42]. Whilst participants were navigating towards a room containing a goal, we did see evidence for a nonspatial representation of context in both hippocampus and OFC, as evidenced by a reliable offset or “separation” between contexts in the neural manifolds for space. However, this effect was less prominent and statistically weaker than the other effects reported here, and did not survive whole-brain correction, and did not persist in either HC or OFC once participants entered the goal room.

Instead, in our study, the most salient way that context influenced neural coding was by compressing spatial codes so that prospective locations signalled by the context lay closer together in neural state space. Thus, when the contextual cue indicated that goals were found on the horizontal axis (H context), the dimension of neural state space representing east-to-west was compressed; similarly, when the cue disclosed that goals lay on the vertical axis (V context), the north-to-south dimension was compressed. Compression effects were visible in a number of brain regions, but they were most prominent in the OFC and hippocampus, where the rooms lying along the relevant dimensions were coded with a highly correlated neural code. Using a spatial encoding model, loosely based on the properties of hippocampal place cells in the rodent, we show that it is possible to elicit compression of this sort via a simple assumption: that both current and prospective locations are encoded jointly in population vectors for allocentric space, i.e., the place code reflects participants’ estimates of both where they are, and where they expect to be. This occurs because jointly encoding prospective locations (the two spatial goals) that lie on a common axis leads to correlated neural signals along this axis, which in turn are visible in compressions of the neural geometry for space.

The compression in HC and OFC was sufficiently prominent that in our context-dependent navigation task, neither region naively reliably encoded the full spatial layout, as might be expected from a pure place code. This might seem curious, given that the hippocampus encodes a spatial representation of the environment in rodents. However, there are a number of possible explanation for this. Firstly, it seems likely that compression is incurred by the need to keep representations of possible spatial plans separate: to ensure that horizontal and vertical goals are not confused. In which case it is possible that compression does not occur in standard navigation paradigms where goals are not flexibly cued from trial to trial, and that there the HC and OFC maps resemble more closely those observed in PPC. Secondly, it is possible that there are variations in the extent to which current and goal locations are decodable from human BOLD signals relative to neuronal recordings in rodents. Indeed, prospective information seems to be a prominent component of human BOLD responses in a variety of settings [9,17,34,53,54], and the place code seen ubiquitously in rodents is much less prominent in monkeys [55] and humans [56]. It is also possible that this is due to differences in recording methods; there is considerable debate about how to jointly understand effects recorded at the micro-, meso- and macroscopic levels during spatial navigation [57].

Our results are thus consistent with the finding that both hippocampus [9,13,34,37,38] and orbitofrontal cortex [40,58] explicitly code for future goal locations, including in human recordings made with both BOLD [34] and intracranial electrodes [13,14]. Our model suggests that the representation of space in the BOLD signal in hippocampus and OFC can be explained by the simple principle that current and prospective (goal) locations are encoded in temporal proximity, but our recording methods do not have the resolution in space or time to detail exactly how that might occur. For example, prospective locations or goals may be represented through dedicated cell types [38] or may be evoked during forwards or backwards simulation occurring via replay mechanisms [59,60], which has also been observed in humans [61]. More generally, our results are consistent with the view that the OFC (and to a lesser extent HC) represents the “task space”, that is it encodes states in a format that is optimised for reward-guided action and planning [62,63]. This could explain the context dependent compression of vertical / horizontal rooms, which represent the reward-relevant axes of our task.

We observed another curious effect by which the neural geometry of the environment was “flipped” between periods in which (i) navigation was ongoing, and (ii) where the goal room had been reached. It is not clear to us what purpose is served by this aspect of the geometry, which was most prominent in HC and OFC. However, it is reminiscent of recent reports that memory traces are rotated in neural state space to prevent them from interfering with perceptual information [64]. In fact, other reports have emphasised a related effect: that retrieval induces spatial memories to be mutually repulsed, perhaps so that they can become more distinguishable [43]. It seems possible that retrieval-based repulsion (between periods) and context-based compression (or attraction) of spatial memories can co-exist; this may be an interesting avenue for future research.

We also examined how the agent’s current location and the location of the navigational goal were jointly represented. Remarkably, we observed that the representation of agent location and goal location is nested, especially in PPC; there is a prominent quadrilateral representation of the currently occupied room, but nested within each room representation is another quadrilateral representation of the current navigational goal. Consistent with the strong effect of period, we see that (in PPC at least) this representation is distorted so that the goal corresponding to the current room is represented distinctly from all other goals. This hierarchical representation of goals and space was not observed in HC or OFC in our study, presumably because averaging over contexts removes the subspace in which location is represented.

How context biases the encoding of sensory signals has been extensively studied in tasks that require a single action to be taken to elicit an outcome, such as visual categorisation. In these tasks, different computational mechanisms have been proposed for preventing interference between different tasks (or goals) that are required in different contexts. For example, when the task requires monkeys to classify stimuli into common groups, single neurons or populations in PFC to code for stimuli associated with a given class [6,65–67], echoing the compression of target information reported here. Other reports, however, argue that during categorisation, neural signals coding for different groups are offset by a one-dimensional signal, giving rise to a neural “separation” similar to that tested here [68,69]. Where there are explicit contextual cues signalling the task, context-irrelevant information can be compressed in BOLD signals [7] and is often coded along a perpendicular dimension in neural state space, for example to avoid catastrophic interference [6,7]. Thus, orthogonalization, separation and compression are candidate mechanisms for mediating the contextual modulation of sensory codes in both instantaneous and sequential decision tasks.

## Acknowledgements

This work was supported by a Sir Henry Wellcome Fellowship (award 210849/Z/18/Z) to P.S.M-K., by generous funding from the European Research Council (ERC Consolidator awards 725937 to C.S. and 820213 to G.P. and ERC Starting grant 852669 to N.W.S.), by Special Grant Agreement No. 945539 (Human Brain Project SGA) to C.S., H.S., and G.P, and by funding from the Max Planck Society (Max Planck Research Group grant M.TN.A.BILD0004) and from the Federal Government of Germany and the Laender under the Excellence Strategy to N.S.

## Methods

### Participants

Thirty-one human participants were recruited for the experiment through the recruitment system at the Max Plank Institute for Human Development (Berlin). One participant was omitted from the analysis due to a neural structural abnormality and another three participants were omitted due to technical difficulties with the MRI equipment. All analyses were performed on the remaining 27 participants (11 male, 16 female; age 27.3 ± 4.4 years). Participants were compensated for their time at a base rate of €10/hour, plus an extra €10 for participating in an MRI experiment, and finally an additional bonus of up to 10€ (5€ per session) depending on their performance. Informed consent was given before the start of the experiment. The study was approved by the Department of Education and Psychology at the Freie Universität Berlin and the Medical Science Inter-Divisional Research Ethics Committee (R49432/RE001) at the University of Oxford.

### Design, task and procedures

The experiment involved two sessions undertaken on different days. In both sessions, participants performed a computerised task that was built and delivered in the Unity 3D games environment. The task involved navigating an avatar through a grid world to collect rewards. On day 1, participants performed a training task outside of the scanner, using the arrow keys on a laptop computer to move the avatar through the environment (see **Fig. 1A**). On day 2 (32.0 ± 3.6 hours later), they performed the task lying supine in an MRI scanner, viewing the screen through a mirror and using an MRI-compatible button box to respond.

On both days, the grid world environment was composed of four adjoining rooms arranged in a square. We refer to the rooms as southwest (SW), northwest (NW), southeast (SE) and northeast (NE) rooms. Each room was composed of 4 x 4 grid squares and was connected to the two cardinally adjacent rooms (e.g., SW was connected to SE and NW but not NE) via a single “bridge” square. It thus mirrored the classic “four rooms” environment commonly used in AI research [70]. At each point in the trial, participants could only see the 4 x 4 squares of the currently occupied room, plus the two additional bridge squares; the other rooms were offscreen. One square of each room contained a boulder, and two of the four boulders in the environment were associated with a reward (the reward was revealed when the avatar collided with the boulder). During training, the grid squares were differently coloured in each of the four rooms (in order to help people learn to navigate); during test, they were all purple. Traversing a bridge square incurred a variable delay during which the full map was briefly shown. This was to encourage participants to consider their room choices carefully before moving between rooms, and later on day 2, to more easily separate the BOLD response pertaining to the occupation of different rooms.

On both days task was divided into blocks of 16 trials (n = 4 during day 1; n = 6 during day 2. In the scanner, these constituted 6 independent scanner runs). On each trial, participants began in the inner corner of a randomly chosen room (they could identify the room by the locations of the visible bridge squares). Before navigation began, participants were shown a contextual cue, which was a picture of one of two food items (**Fig. 1C**). Unbeknownst to participants, each cue disclosed one reward location conditional on the other for that trial. For example, in scanner run 1, cue A (a martini icon) indicated that the rewards were in rooms lying in the same horizontal axis, and cue B (a peanut icon) that the rewards were in the same vertical axis (with neither disclosing which specific rooms). Different pairs of food items were chosen on day 1, and then on blocks 1-2, 3-4, and 5-6 of day 2 (4 pairs total) so that participants had to generalise the structure to previously unseen items. Food icons used for the day 2 (scanner) task included watermelon, cheese, mushroom, avocado, pineapple and banana.

Participants navigated freely (up, down, left, right) using buttons, causing the avatar to move within the environment (the background grid remaining fixed). When participants alighted on a boulder that was associated with a reward, the reward was revealed by showing the food item that had been cued on that trial, before navigation could recommence. Participants were instructed to find the two rewards as quickly as possible and received a *trial score* that was equal to the number of seconds the participant had remaining on their timer at the end of each trial. If participants took longer than the timer deadline to find both rewards, 20 points were deducted from their total trial score. The task was calibrated so that a participant that ignored the cues and navigated to boulders in any order would only meet the deadline on approximately 50% of trials, set to be 40s on day 1 and 50s in the scanner on day 2. Aggregate trial score was converted to a financial bonus at the end of the experiment.

The timing of events within each trial were as follows. Each trial started with the controls disabled, and the location of the avatar in the start room was shown for 2.5s. The contextual cue was then displayed enlarged in the centre of the screen for 1.5s. After a further 1s the controls were enabled, and participants were able to move the avatar through the environment by pressing arrow keys (day 1) or button box keys (day 2). At this point, the timer (visible in the top right hand corner of the screen) started ticking down from a deadline value (40s on day 1, 50s on day 2). On day 2, participants could move the avatar at a maximum speed of 1 grid square every 0.4s (increased from 0.25s on day 1, to ensure participants remained in each room long enough to obtain a clear per-room neural signal). When moving through a hallway, controls were disabled for a period of time before players were able to move again, where this period was drawn from a truncated exponential distribution (mean 2s; min 1.5s; max 7s). When the avatar collided with a boulder, the controls were again disabled for a period (sampled from truncated exponential with mean 2s; min 1s; max 5s) while either a reward or no reward was shown. At the end of the trial a message saying “well done” appeared on the screen and the participant’s total score was visibly updated using the remaining seconds left on the timer, which corresponded to additional points. After each trial participants were shown a black screen for a period before the next trial began (ITI sampled from truncated exponential distribution with mean 2.5s, min 1.5s, max 7s).

In our design, we consider the cues to be “contexts” signalling whether rewards were found on the horizontal or vertical axes of the four rooms environment. However, because rewards could be in any room (i.e., in a vertical condition they could be in the SW and NW or the SE and NE), occupancy probabilities were closely matched across contexts. Nevertheless, to ensure good coverage of the environment, and to attempt to match transitions as well as occupancy, we also introduced a “robot control” phase in every trial, in which either the robot or the participant began by controlling the avatar. If the robot controlled the avatar, it moved at approximately the same pace as an average participant, and typically to a non-rewarded room, where it made a beeline for the boulder. Every time the computer (participant) reached a boulder, the control was passed back to the participant (computer) until the next boulder was reached. On average, the amount of time spent per trial under control of the robot was 11.7 seconds, compared to 14.3 seconds under participant control. We also verified that the neural representation of space independently for the human- and robot-controlled phase (**Fig. S8**).

The behavioural training session (day 1) began with two practice trials, which used different contextual cues, which did not signal the location of one cue conditional on the other. During the training session (day 1), cues were blocked so that participants alternated between horizontal and vertical contexts in an ABAB design. During day 2 in the scanner, contexts were interleaved from trial to trial, so that different contexts were not associated with distinct, prolonged temporal episodes.

Each run contained 16 trials, which were balanced across pairs of runs with the same reward cues. Trials were balanced across the two cues (32 trials / 2 cues = 16), starting rooms (16 trials per cue / 4 rooms = 4), whether the start room was rewarded or not (4 trials per room per cue / 2 = 2), and whether the participant foraged first or the robot foraged first (2 trials rewarded per room per cue / 2 = 1). Trial ordering was randomised across participants.

At the end of the scanning sessions participants responded to a situational quiz which examined their explicit understanding of the reward covariance rules. They were asked four questions of the form “*You have just found a cheese in the top right room. Which room will the other cheese be in?*”, and four questions which tested their counterfactual understanding, such as “*You were looking for a cheese and did NOT find one in the bottom right room. Which rooms will contain the two cheeses?*”. Each participant was assessed with a version of the quiz that mentioned the reward pair that they had most recently observed (those in the final two runs). The maximum possible score was 8. We examined the correlation across participants between these quiz scores and (1) the average first room choice accuracy in the scanning session, and (2) the mean trial score in the scanning session.

### Behavioural analysis

We computed three behavioural metrics. Firstly, we considered the trial score, which is roughly proportional to the average time taken to complete a trial. Secondly, we computed the *transition bias*, which is the relative fraction of transitions between horizontal and vertical rooms in the appropriate context (H or V), computed across both human- and computer-controlled events:

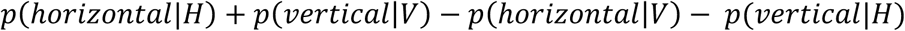

Thirdly, we computed first choice accuracy, which is the probability that participants made an optimal transition from the first room occupied to the second. This measure is particularly sensitive because a participant who perfectly understand the meaning of the contextual cues can always head to the correct 2^nd^ room, on the basis of whether the 1^st^ (starting) room contains a reward or not. First choice accuracy was quite highly correlated with transition bias (r = 0.86).

### Computational model

We defined a simulated environment corresponding to the four rooms arena, in which locations were denoted by values in the range [-1,1] in both the *x* and the *y* dimension. We rescaled participants’ observed movement trajectories through the grid environment so that they mapped onto this simulated environment and located the boulders at their approximately corresponding positions. This allowed us to model neural responses using an encoding model that consisted of simulated place cells. The place cells exhibited bivariate Gaussian response fields that regularly tiled the arena on a 10 × 10 square lattice (but the results we describe were very similar, albeit but more variable, if we drew their tuning preferences from random uniform distributions; we also verified that almost identical results are obtained if we truncate place fields so that they do not straddle different rooms). We assume that there are 200 place cells in each context (two cells coding for each location).

In this simple model, we define the place field of neuron *i* in context *c* as peaking at an [*x*, *y*] location *θ_i_*(*c*). We note that the vector of place fields in the two contexts *θ*(*c* = *V*) and *θ*(*c* = *H*) may be the same, or partially or fully different. This is controlled by the parameter *β*, which determines the fraction of cells for which *θ*(*c* = *H*) ≠ *θ*(*c* = *V*). Thus if *β* = 0 then all cells code for the same location regardless of context (no remapping), if *β* = 0.5 then 50/100 cells exhibit overlapping place fields between contexts (partial remapping), and if *β* = 1 then all cells change their tuning between contexts (full remapping).

Thus, in any given context we can estimate the neural response of neuron *i* on time step *t* as being

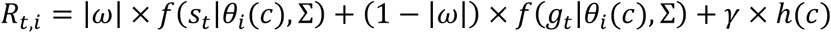

In the expression above, *f*(·|*θ_i_*, *σ*) is the bivariate normal distribution evaluated at the preferred [*x*, *y*] tuning location (place field) for neuron *i* in context *c*, and Σ = [0.25, 0; 0 0.25]. We define the current [*x*, *y*] location of the agent as *s_t_*, whereas *g_t_* is the [x, y] coordinate location of the goal to which the agent is headed on the current timestep. Finally, ℎ(*c*) is a context-specific neural signal, which depends uniquely on the current context (H or V) active on that timestep and not on the location of the agent or goal.

In addition to orthogonalization (controlled by *β*), separation and compression are controlled via two further free parameters. The gain parameter *γ* determines the relative influence of ℎ(*c*), the (place-insensitive) signal coding for context, which has the effect of neural separation between neural manifolds for space in each context. Finally, the mixing parameter *ω* determines the relative influence of the current (*s_t_*) and prospective (*g_t_*) location on the neural population response. Where *ω* = 0, only the participant’s current location is encoded, as in a “classical” place field model. Where *ω* > 0, the model encodes a mixture of the current location and the prospective (goal) location. We also allow for *ω* < 0; in this case, we use the closely related expression

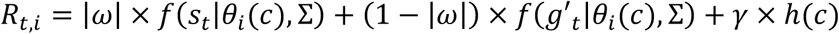

Where *g*′*_t_* is a fictitious goal location that is swapped on the horizontal / vertical axis, as if participants were prospectively encoding horizontal locations in the V conditions and vertical locations in the H condition. This entails that space is compressed along the dimension perpendicular to the axis on which the two goals can be found. We call this anti-compression.

We use the observed individual trajectories, and the actual prospective goals (i.e., where participants were genuinely headed) at each time point to evaluate the model for each agent. This provides us with a neural response matrix of size 200 × *t* in each of 48 trials in context H and 48 trials in context V. We then average those timepoints in which the avatar was in the SW, NW, SE and NE rooms for each context, yielding a 200 × 8 matrix, which we use to generate an 8 × 8 RDM (expressing correlation distance) for each simulated participant. For visualisation, we average these RDMs across the simulated cohort, and plot them using multidimensional scaling (e.g., **Fig. 2C**).

### fMRI data collection and pre-processing

#### Anatomical MRI data

MRI data were acquired at the Max Planck Institute for Human Development in Berlin using a 32-channel head coil on a 3T Siemens Magnetom Triotrim MRI scanner (Siemens, Erlangen, Germany). At the start of the scanning session, a T1-weighted (T1w) high-resolution anatomical image was obtained using a Magnetization Prepared Rapid Gradient Echo (MPRAGE) sequences (sequence parameters: repetition time (TR) = 2500 ms, echo time (TE) = 4.77 ms, flip angle = 7°, field of view (FOV) = 256 mm; voxel size = 1 x 1 x 1 mm).

Data were processed within the fMRIPrep framework. The T1w image was corrected for intensity non-uniformity with N4BiasFieldCorrection [71], distributed with ANTs 2.2.0, and used as T1w-reference throughout the workflow. The T1w-reference was then skull-stripped with a Nipype implementation of the antsBrainExtraction.sh workflow (from ANTs), using OASIS30ANTs as target template. Brain tissue segmentation of cerebrospinal fluid, white matter and gray matter was performed on the brain-extracted T1w using fast (FSL 5.0.9). Brain surfaces were reconstructed using recon-all (FreeSurfer 6.0.1), and the brain mask estimated previously was refined with a custom variation of the method to reconcile ANTs-derived and FreeSurfer-derived segmentations of the cortical gray matter of Mindboggle [72]. Volume-based spatial normalization to MNI space (MNI152NLin2009cAsym) was performed through nonlinear registration with antsRegistration (ANTs 2.2.0), using brain-extracted versions of both T1w reference and the T1w template.

#### Functional MRI data

Functional MRI data were acquired using a T-2-weighted (T2w) echo planar imaging (EPI) pulse sequence sensitive to BOLD contrast (sequences parameters: TR = 2000 ms, TE = 30 ms, FOV = 192 x 192 mm, flip angle = 80°, voxel size = 3 x 3 x 3 mm). The task was divided into 6 functional runs, each lasting between 10 and 15 minutes, depending on participant performance.

For each of the six scanning runs, the following pre-processing steps were performed. Initially, a reference volume and its skull-stripped version were generated using a custom methodology of fMRIPrep. A B0-nonuniformity map was estimated based on two EPI references with opposing phase-encoding directions, with 3dQwarp (AFNI 20160207). Based on the estimated susceptibility distortion, a corrected EPI reference was calculated for a more accurate co-registration with the anatomical reference. The BOLD reference was then co-registered to the T1w reference using bbregister (FreeSurfer) which implements boundary-based registration [73]. Co-registration was configured with six degrees of freedom. Head-motion parameters with respect to the BOLD reference (transformation matrices, and six corresponding rotation and translation parameters) are estimated before any spatiotemporal filtering using mcflirt (FSL 5.0.9). BOLD runs were slice-time corrected using 3dTshift from AFNI 20160207 and the time-series were resampled to their original native space as well as to the standard MNI space. BOLD data were moreover smoothed with a 6 mm full-width half-maximum Gaussian kernel. A reference volume and its skull-stripped version were generated using a custom methodology of fMRIPrep. Several confounding time-series were calculated based on the pre-processed BOLD: framewise displacement, DVARS and three region-wise global signals. FD and DVARS are calculated for each functional run, both using their implementations in Nipype (following the definitions by [74]).

### fMRI analysis: first level GLMs and ROI definition

The pre-processed BOLD timeseries data were modelled with general linear models (GLMs) that contained regressors for different task events. The first GLM (GLM1) contained the following regressors: the contextual cue, the movement period of each trial before the first feedback when subjects had no knowledge about the locations of the reward, the subsequent movement periods when the agent was occupying a room without a reward (pre-goal room period), the subsequent movement periods when the agent was occupying a room with a reward (goal room period), those periods when the agent occupied a hallway between adjacent rooms, the feedback periods when a reward was presented, and the feedback periods when no reward was presented. Note that all movement periods after the first feedback were labelled in a consistent manner, based on the presence vs. absence of reward in the current room. Hence, the second movement period of each trial (i.e., when the agent is still in the first room after receiving the first feedback) would be treated as pre-goal room or goal room, depending on the outcome of the first feedback.

We modelled data from both the self-directed and robot control periods together to ensure adequate coverage of space in both contexts (see Figure S8 for a control analysis showing aligned spatial representations during both types of events). Note that we defined separate regressors for pre-goal room periods and goal room periods for each of the four rooms of the grid world (SW, NW, NE, SE) and each behavioural context (vertical vs. horizontal goal alignment), resulting in 16 regressors for movement periods after the first feedback. The GLM also contained nuisance regressors pertaining to participants’ head motion (three rotation parameters and three translation parameters), the global signal in the white matter, and the framewise displacement. All regressors were modelled with variable durations from start to finish and convolved with a canonical haemodynamic response function. To ensure sufficient trial counts, we concatenated the BOLD time-series data across odd and even scanning runs (using the spm_fmri_concatenate function). We conducted a second GLM (GLM2) that defined movement periods in terms of the current location of the agent (SW, NW, NE, SE) and the prospective location of the navigational goal (SW, NW, NE, SE). This GLM collapsed events across behavioural contexts (i.e., trials with vertical vs. horizontal goal alignment), thereby also resulting in 16 regressors for movement periods after the first feedback.

Five regions-of-interest (ROIs) were defined based on existing atlases. We used the Wake Forest University Pickatlas (integrated into SPM) to define ROIs for the hippocampus (bilateral areas labelled *Hippocampus*), orbitofrontal cortex (bilateral areas labelled *Frontal_Inf_Orb*; *Frontal_Mid_Orb*, *Frontal_Sup_Orb*), and visual cortex (bilateral areas labelled *Occipital_Mid*). ROIs for prefrontal and posterior parietal cortices were defined based on an atlas provided by Fedorenko et al. (2013) that delineates frontoparietal brain areas implicated in cognitive control across a variety of cognitive domains (the whole atlas is available for download at http://imaging.mrc-cbu.cam.ac.uk/imaging/MDsystem).

### fMRI analysis: neural geometry

To compute the neural geometry, we obtained the multivariate pattern evoked by each of the 16 predictors (in either GLM1 or GLM2) for each participant in each scanner run. Thus, for each region (or searchlight) this yielded a *n*(*v_r_*) × 16 × 6 data array, where *n*(*v_r_*) is the number of voxels in region *r*. We collapsed over odd and even runs, giving us two *n*(*v_r_*) × 16 arrays. Using a previously described method called ‘reliability-based voxel selection’ [75], we began by identifying eligible voxels for multivariate analysis (feature selection). We correlated, in each individual voxel, the pattern of activity over the 16 conditions between odd and even runs. Voxels with Pearson’s *r* > 0 were included in all multivariate analysis. This left a minimum of 1470, 1931, 709, 320 and 1335 voxels in visual, PPC, PFC, hippocampus and OFC ROIs respectively. This feature selection method ensures that only those voxels with consistent patterns across runs (i.e., those with higher signal) are included in the analysis (but it does not specify what the pattern should be in these voxels). To verify that this did not bias our analysis in any way, we reran all analyses in the paper on shuffled data to which we applied the same feature selection methods. To achieve this, we shuffled the mapping across voxels independently between training and test, creating a dataset with equivalent summary statistics but no train-test consistency, and reran our analyses including the feature selection stage. We observed no deviation from the expected null distribution in this case. Next, we used singular value decomposition to reduce the dimensionality to *d* dimension within the ROI; for all statistical analyses, we used *d* = 10 (we did not apply this step for MDS visualisation). The first 10 principal components captured about ∼60% of the variance in most regions and participants

At this stage (where required, i.e., for **Fig. 3** and **Fig. 4**) we separated the ROIs into pre-goal room and goal room periods. This was desirable because of the large offset in behaviour between these periods, but we obtain very similar results when we analyse all the data together (**Fig. 5**). In each case, we computed RDMs (8 × 8 or 16 × 16) in cross-validation; this means that we computed *RDM*_*ij*_ which is the dissimilarity between the neural pattern from the *i*^*th*^ condition in odd scanner runs to the *j*^*th*^ condition in even scanner runs. We then averaged these RDMs about the diagonal and regressed lower triangle of the RDM against that of a predictor matrix composed of one or more (standardised) model RDMs, obtaining beta coefficients for their competitive fit. The diagonal is not zero for our data RDMs because of the cross-validation step, so we set it to zero for MDS visualisation only; note that this has no impact on our statistical analysis. The RDMs we generated were designed to be as orthogonal as possible. For example, the “map” RDM captures similarity structure over and above that in the “room” RDM by predicting larger distances on the diagonals (e.g. NE to SW) than edges (e.g. NE to NW; see **Fig. 3E**. The median Pearson correlation between predictors was r = .0.175 and no pair of predictors had a Pearson’s correlation that exceeded 0.5.

For the “scores” analysis, we generated three perfectly orthogonal matrices that we call *compression*, *separation* and *map*. Each of these score matrices is the same size as the RDM and comprises binary values (+1 and -1) for key condition pairs; each sums to zero. Each score matrix is multiplied elementwise with the data RDM for each participant and averaged, yielding a score that is > 0 if matrix values set to 1 are more dissimilar than those set to -1 and < 0 otherwise. This allows us to do group-level one-sample t-tests (against zero) to test for these three effects.

The *compression* matrix tests whether the east and west rooms are more similar in the H condition, and the north and south rooms in the V condition, than the converse (it is thus the subtraction of the compression and anti-compression matrices shown in **Fig. 3E**). The *separation* matrix sets values between contexts to +1 and those within contexts to -1, excluding the minor diagonals (i.e., the dissimilarity between each room and itself across contexts). The *map* matrix tests whether each plane is shaped like a quadrilateral, mirroring the geometry of the four rooms environment. To this end, the scores matrix has values of +1 for the diagonals (e.g., SE to NW) and -1 for the cardinals (e.g., SE to NE; this is done in subsets to ensure the matrix sums to zero). The resulting vector of scores across the participant cohort is correlated with behavioural measures, including first choice accuracy and transition bias, using Pearson’s correlation.

In the angle analysis, we assess the angle between the north-south and east-west vectors within contexts, across contexts, and across contexts and goal room period. We do this using the full 16 × 16 matrix. We first compute the difference in high-dimensional vector coding for each room in each context and period, leading to a data matrix of size *n*(*v_r_*) × 16 × 16. We manually compute the angle between those edges predicted to be parallel, orthogonal or inverted in the model in **Fig. 5C** and plot these in **Fig. 5D**.

To detect potential signals outside of the chosen ROIs, we repeated the “scores” analysis, described above, in a whole-brain searchlight approach, where RSA was conducted at each voxel with a group of surrounding voxels (spherical searchlight radius = 12 mm). Analogously to the ROI analyses, we conducted separate analyses for pre-goal room period and goal room period. For each voxel within the searchlight, we extracted the 8 beta coefficients from the GLM corresponding to the regressors for each room (SW, NW, NE, SE) and context (vertical, horizontal). We next applied voxel selection and dimensionality reduction, as described above, and computed cross-validated RDMs (8 x 8), which were multiplied elementwise with each predictor matrix, yielding three whole-brain maps with regression coefficients for each subject. These maps were smoothed using an 8 mm FWHM Gaussian kernel. Statistical significance was established separately for each voxel by testing the regression coefficients against zero using one-sample t-tests. Correction for multiple comparison was conducted via family-wise error correction (p < 0.05) as implemented in SPM 12.

