## Supplementary Materials for "Goal-seeking compresses neural codes for space in the human hippocampus and orbitofrontal cortex"

Supplementary Figure 1

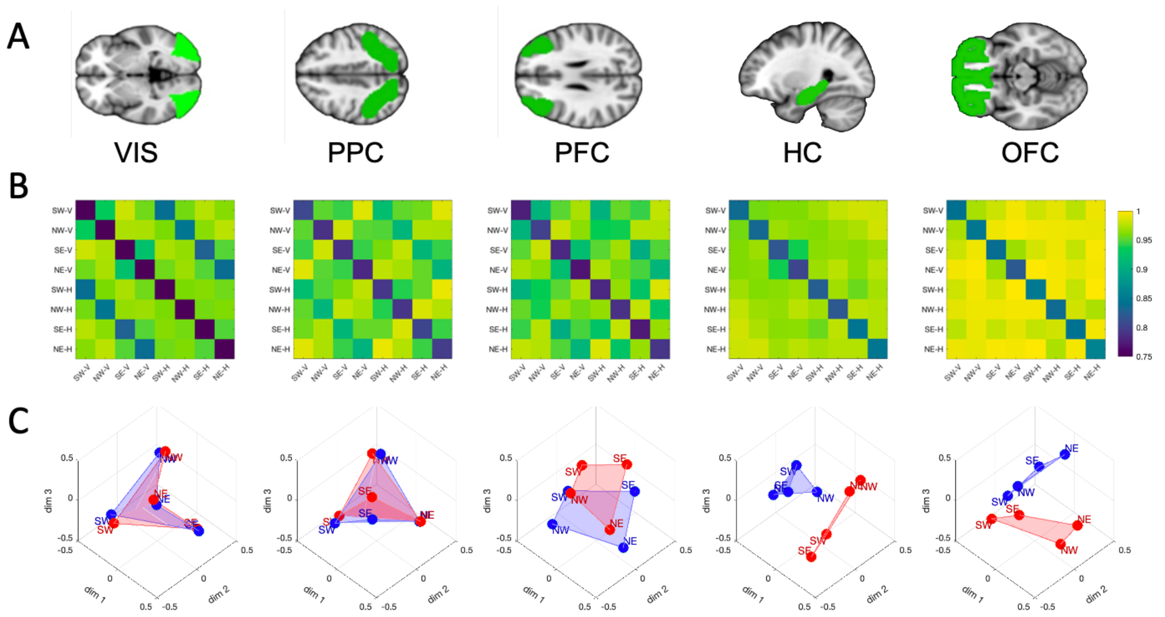


**Figure S1. RDMs and neural geometry from the goal room period. (A)** Regions of interest, shown again for convenience **(B)** Group average RDMs for each ROI. Each 8 × 8 RDM is ordered {SW,NW,SE,NE} for first the vertical and then the horizontal context. Warmer colours indicate greater dissimilarity, and cooler colours greater similarity. **(C)** MDS plots (from the group average RDM) for each region. Blue dots are rooms in the vertical context and red in the horizontal context. For legibility, cardinally adjacent rooms within a context are linked by lines, which collectively form a quadrilateral when allocentric space is coded in just 2 dimensions.

Supplementary Figure 2

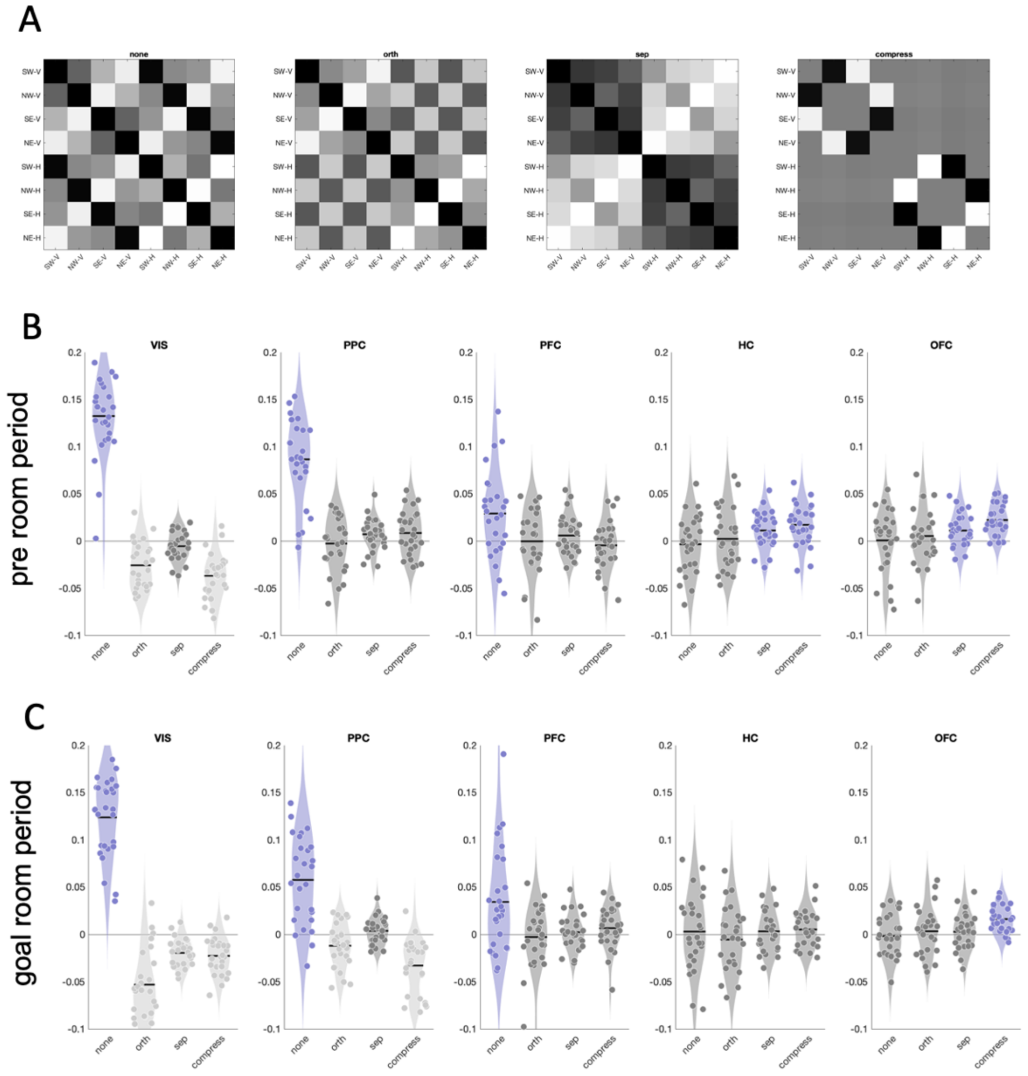


**Figure S2. Correlations with RDMs from the place field model. (A)** RDMs generated from the best fitting variant of the place field model, under parameterisations where (i) no parameters were allowed to vary ("none"); (ii) only the orthogonalization ($\beta$) parameter is allowed to vary; (iii) only the separation ($\gamma$) parameter is allowed to vary; and (iv) only the compression ($\omega$) parameter is allowed to vary. Lighter colours indicate greater dissimilarity. **(B)** Coefficients from a regression on the data RDM for each region, from the pre-goal room period. **(C)** same as (B) but for the goal room period.

Supplementary Figure 3

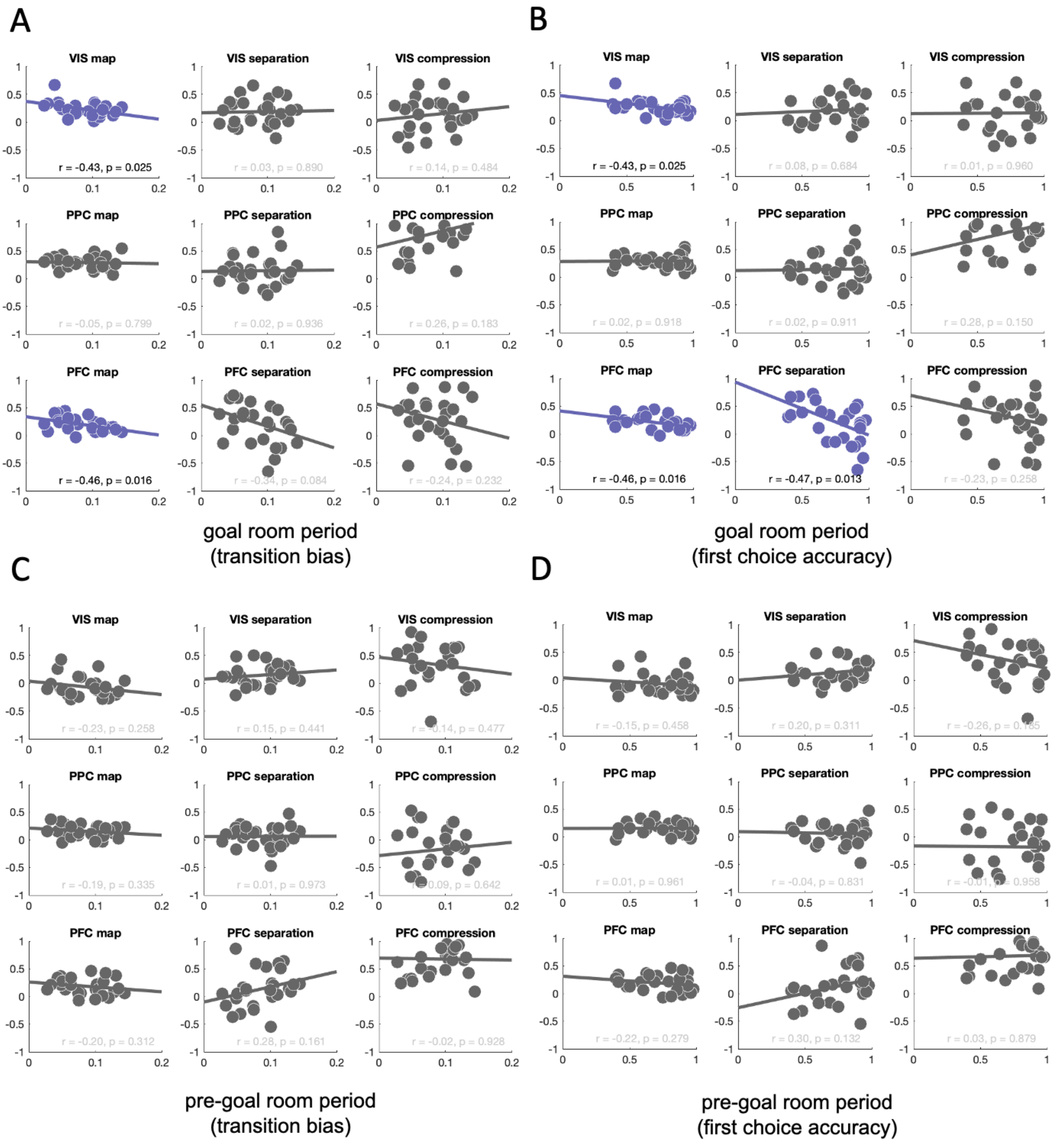


**Figure S3. Correlations between neural scores (map, separation and compression) and behaviour for visual cortex, PPC and PFC.** (A) Correlations with transition bias for the goal room period; (B) Correlations with first choice accuracy for the goal room period; (C) Correlations with transition bias for the pre-goal room period; (D) Correlations with first choice accuracy for the pre-goal room period. Each dot is a single participant, and the line is the best linear fit. Blue colouring is used to highlight significant correlations ($p < 0.05$)

Supplementary Figure 4

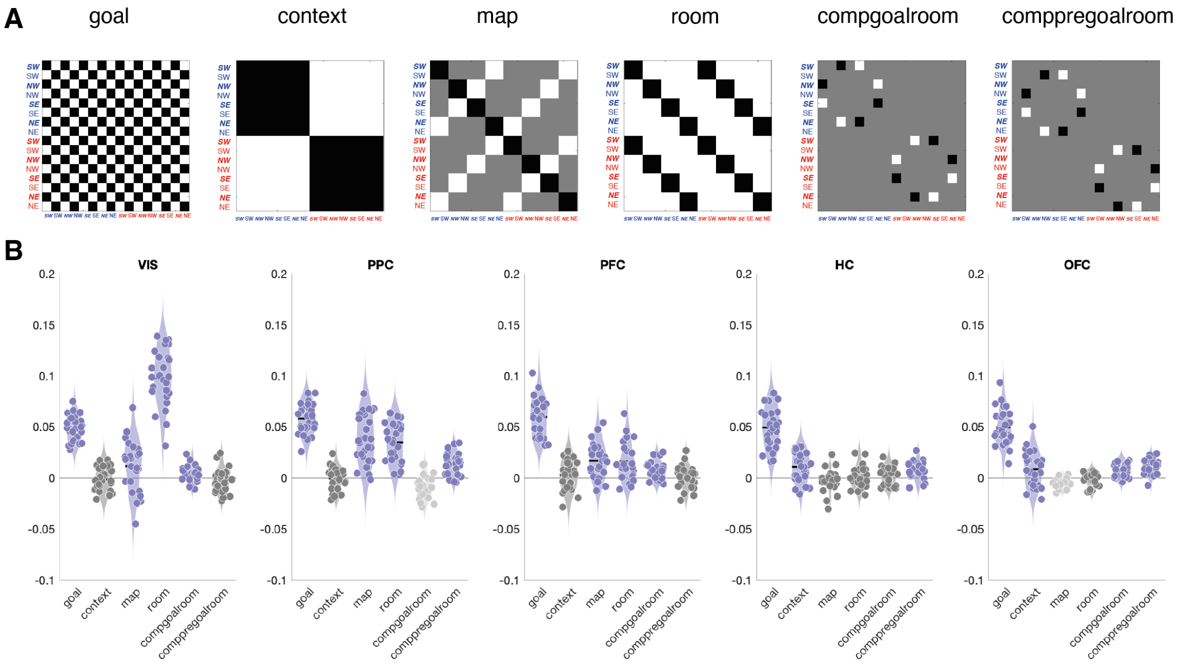


**Figure S4. (A)** model RDMs used for the analysis shown in Fig. 5. **(B)** Coefficients for the regression of model RDMs for the full 16 × 16 (period × room × context) analysis described in Fig. 5. Blue dots show significant ($p < 0.01$) predictors.

Supplementary Figure S5:

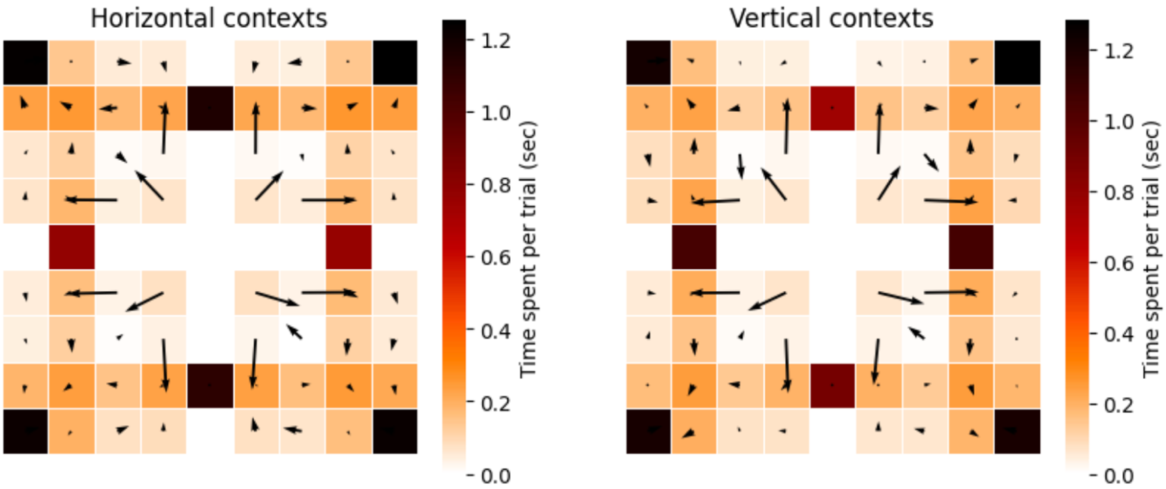


**Figure S5.** Heatmaps of the average grid square occupancy per trial in each of the two contexts for human-controlled movement periods only. Black arrows show the average transition vector from each grid square. Data are averaged across participants.

Supplementary Figure 6:

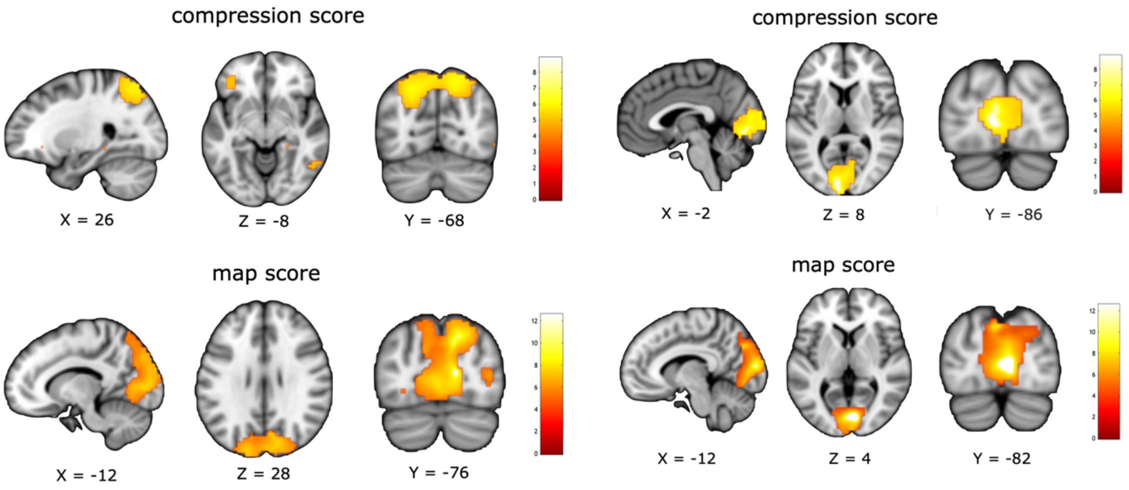


**Figure S6.** Searchlight analyses: whole-brain effects of *compression score* and *map score* for the pre-goal room period (left) and the goal-rom period (right), rendered onto a template brain after familywise error correction at $p < 0.05$.

Supplementary Figure 7:

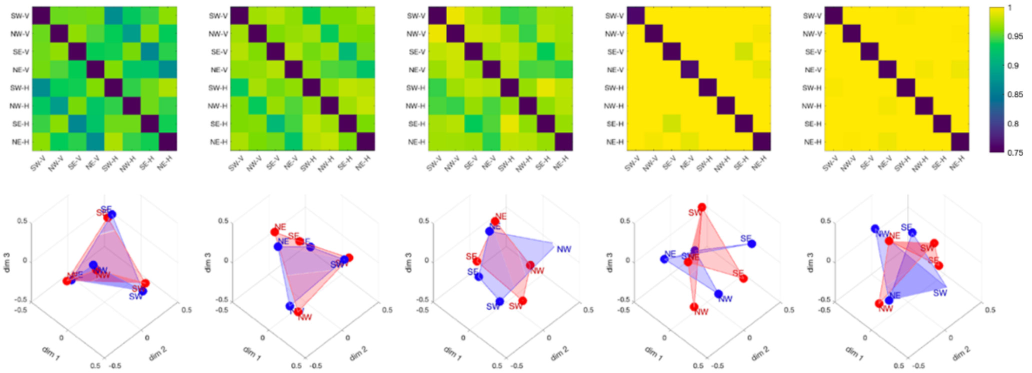


**Figure S7.** Neural geometries from an analysis focusing on brain activity during the first movement period of the trial (i.e., before any feedback has been received and agents have no knowledge of reward locations). The top row displays group average RDMs for each ROI. Each 8 × 8 RDM is ordered by room {SW,NW,SE,NE} for first the vertical and then the horizontal context. Warmer colours indicate greater dissimilarity, and cooler colours greater similarity. The middle row displays MDS plots (from the group average RDM) for each region. Blue dots are rooms in the vertical context and red in the horizontal context. For legibility, cardinally adjacent rooms within a context are linked by lines, which collectively form a quadrilateral when allocentric space is coded in just 2 dimensions.

Supplementary Figure 8:

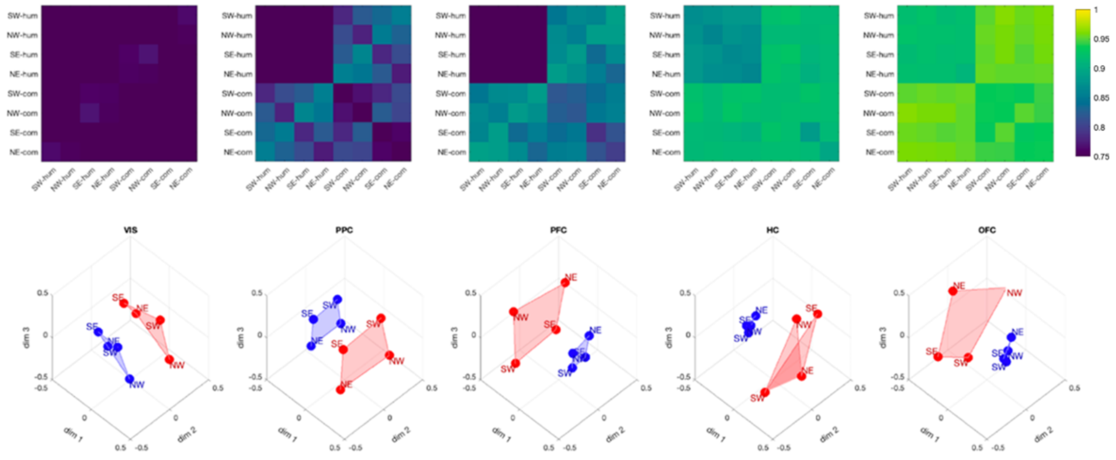


**Figure S8.** Neural geometries from an analysis focusing on brain activity during movement periods after the first feedback, with separate predictors for events that were controlled by the human participant and the computer. The top row displays group average RDMs for each ROI. Each 8 × 8 RDM is ordered by room {SW,NW,SE,NE} for first the human and then the computer controlled events. Warmer colours indicate greater dissimilarity, and cooler colours greater similarity. The bottom row displays MDS plots (from the group average RDM) for each region. Blue dots are rooms in the human-controlled events and red dots are rooms in the computer-controlled events. For legibility, cardinally adjacent rooms within a context are linked by lines, which collectively form a quadrilateral when allocentric space is coded in just 2 dimensions. **x**
